# Sanglifehrin A mitigates multi-organ fibrosis in vivo by inducing secretion of the collagen chaperone cyclophilin B

**DOI:** 10.1101/2023.03.09.531890

**Authors:** Hope A. Flaxman, Maria-Anna Chrysovergi, Hongwei Han, Farah Kabir, Rachael T. Lister, Chia-Fu Chang, Katharine E. Black, David Lagares, Christina M. Woo

## Abstract

Pathological deposition and crosslinking of collagen type I by activated myofibroblasts drives progressive tissue fibrosis. Therapies that inhibit collagen synthesis by myofibroblasts have clinical potential as anti-fibrotic agents. Lysine hydroxylation by the prolyl-3-hydroxylase complex, comprised of cartilage associated protein, prolyl 3-hydroxylase 1, and cyclophilin B, is essential for collagen type I crosslinking and formation of stable fibers. Here, we identify the collagen chaperone cyclophilin B as a major cellular target of the macrocyclic natural product sanglifehrin A (SfA) using photo-affinity labeling and chemical proteomics. Our studies reveal a unique mechanism of action in which SfA binding to cyclophilin B in the endoplasmic reticulum (ER) induces the secretion of cyclophilin B to the extracellular space, preventing TGF-β1–activated myofibroblasts from synthesizing collagen type I *in vitro* without inhibiting collagen type I mRNA transcription or inducing ER stress. In addition, SfA prevents collagen type I secretion without affecting myofibroblast contractility or TGF-β1 signaling. *In vivo,* we provide chemical, molecular, functional, and translational evidence that SfA mitigates the development of lung and skin fibrosis in mouse models by inducing cyclophilin B secretion, thereby inhibiting collagen synthesis from fibrotic fibroblasts *in vivo*. Consistent with these findings in preclinical models, SfA reduces collagen type I secretion from fibrotic human lung fibroblasts and precision cut lung slices from patients with idiopathic pulmonary fibrosis, a fatal fibrotic lung disease with limited therapeutic options. Our results identify the primary liganded target of SfA in cells, the collagen chaperone cyclophilin B, as a new mechanistic target for the treatment of organ fibrosis.

## Introduction

Fibrosis is a pathological process characterized by excessive deposition of collagen-rich extracellular matrix in response to chronic or overwhelming tissue injury,^1^ ultimately leading to the development of fibrotic diseases that can affect nearly every organ including the skin,^2^ lungs,^3^ liver,^4^ and kidneys.^5,6^ Progressive tissue fibrosis eventually results in failure of the affected organs and death. Skin and lung fibrosis are hallmarks of fatal fibrotic diseases such as systemic sclerosis (SSc), an autoimmune multi-organ fibrotic disease,^7,8^ and idiopathic pulmonary fibrosis (IPF), an age-related interstitial lung disease.^9^ In these diseases, tissue fibrogenesis is driven by chronic epithelial and vascular damage, type 2 inflammation, and activation of scar-forming cells known as myofibroblasts.^10,11^ Injury-activated epithelial, vascular, and immune cells release pro-fibrotic mediators that promote the activation, differentiation, and survival of myofibroblasts.^8,12,13^ Targeting inflammation with recombinant interferon gamma,^14^ TNF-α neutralizing antibodies,^15^ and immunosuppressive low doses of prednisone and azathioprine^16^ has not shown therapeutic efficacy and may even worsen lung fibrosis in patients with SSc and IPF. Separately, efforts to understand the biology of myofibroblasts have led to the identification of two molecules, pirfenidone and nintedanib, which prevent collagen synthesis induced by the pro-fibrotic cytokine transforming growth factor β1 (TGF-β1).^17,18^ In 2014, the FDA approved these two drugs for the treatment of IPF, marking a turning point for treatment of fibrosis.^19^ More recently, nintedanib has also received approval for the treatment of chronic fibrosing interstitial lung diseases (ILDs) and ILD associated with SSc (SSc-ILD).^20,21^ Despite being approved for clinical use, the mechanisms of action of pirfenidone and nintedanib remain incompletely understood. Pirfenidone has been shown to inhibit TGF-β1–induced collagen secretion by fibrotic fibroblasts through unclear mechanisms,^17^ while nintedanib is a potent multi-kinase inhibitor that acts on platelet-derived growth factor, vascular endothelial growth factor, and fibroblast growth factor signaling.^18^ A greater understanding of these drugs’ molecular target(s) and mechanism(s) of action may allow for improvements upon the modest efficacy and low tolerability due to off-target side effects observed in the current generation of therapies. There is a continuing need to better elucidate the biology of myofibroblasts in order to develop anti-fibrotic agents targeting myofibroblast activation and collagen synthesis with greater selectivity and improved efficacy ^1^.

Advances in our understanding of the biology of myofibroblast activation have led to the identification of novel pro-fibrotic mediators that promote the synthesis, deposition, and remodeling of the extracellular matrix (ECM) in fibrotic disease.^1,5,7,8^ These efforts have enabled the development of both small molecules and biologics aimed at targeting these pro-fibrotic pathways, several of which are currently being tested in phase II and III clinical trials.^22,23^ Novel approaches including drug repurposing are an attractive alternative to de novo development of anti-fibrotic therapies due to the use of de-risked compounds, some of them already tested in clinical trials, potentially shortening development timelines. One of the challenges related to this strategy is the identification of specific targets blocked by agents with anti-fibrotic properties, as exemplified by pirfenidone, whose mechanism of action remains to be determined. Chemical proteomics methods like photo-affinity labeling (PAL) have allowed for unbiased target identification studies across the proteome to illuminate binding interactions inside live cells. In a PAL experiment, a small molecule probe functionalized with a photo-activatable group, such as a diazirine, is added to cells and covalently conjugated to interacting protein targets upon UV irradiation. The labeled proteins are enriched and identified by mass spectrometry (MS).^24,25^ We and others have identified novel protein interactions of small molecule fragments, metabolites, and drug compounds using this approach.^24,26,27^ However, despite the wide use of PAL to revealed novel targets of small molecules in cells, it has yet to be applied to the characterization of novel anti-fibrotic targets.

Here, we apply chemical proteomics to understand the mechanism of action of the natural product sanglifehrin A (SfA), a molecule with anti-proliferative and immunosuppressant properties that was discovered by Novartis in 1999 and whose molecular targets remain only partially understood. SfA is composed of a 22-membered macrocyclic ring decorated with a unique spirolactam that was identified in a screen for bacterially produced compounds that bind to the peptidyl-prolyl *cis*-*trans* isomerase cyclophilin A (PPIA).^28,29^ PPIA is also the target of the immunosuppressive drug cyclosporin A (CsA), and the PPIA:CsA complex binds to and inhibit the phosphatase calcineurin, ultimately blocking the proliferation of activated T cells.^30-33^ Prior investigations have shown that SfA is mechanistically distinct from other immunosuppressants, including CsA and rapamycin,^34^ and that SfA blocks cell proliferation at the G1–S phase transition by induction of p53 expression via NFκB signaling.^35,36^ Recent studies have shown that SfA forms a ternary complex with PPIA and the cystathionine beta synthase domain of inosine monophosphate dehydrogenase 2 (IMPDH2), which contributes to the anti-proliferative effects of SfA, ^37^ and our recent structure–activity relationship studies suggest that SfA may exert its effects through additional targets.^38^ While studies have shown that SfA engages several cyclophilins by *in vitro* biochemical assays,^29^ an accurate profile of the interactions of SfA across all cyclophilins in live cells is challenging, where contributions from both binding affinities and the subcellular localization of SfA and its target proteins contribute to the interaction landscape. We used a chemical proteomics approach in immortalized immune cell lines to discover that SfA primarily interacts with cyclophilin B (PPIB) in live cells. PPIB is an endoplasmic reticulum (ER)–resident peptidyl-prolyl *cis-trans* isomerase^39,40^ that catalyzes the rate-limiting step in collagen folding as part of the collagen prolyl 3-hydroxylation complex, leading us to hypothesize that SfA has anti-fibrotic effects by blocking collagen synthesis in fibrogenic fibroblasts.^41-43^ Our studies reveal a novel mechanism in fibroblasts in which SfA binding to PPIB induces the secretion of PPIB from the ER to the extracellular space, preventing collagen folding and synthesis by TGF-β1–activated myofibroblasts and fibrotic fibroblasts isolated from patients with IPF. We provide chemical, molecular, functional, and translational evidence that this mechanism plays an important role in collagen synthesis by pro-fibrotic myofibroblasts *in vitro* and in the development of the lung and skin fibrosis *in vivo* in a preclinical mouse model. We further show that SfA reduces collagen type I secretion by tissue from patients with IPF. Together, our findings identify PPIB as a major cellular target of SfA and demonstrate the therapeutic potential of inhibiting collagen type I synthesis by depletion of intracellular PPIB in fibrotic diseases such as SSc and IPF.

## Results

### Identification of cyclophilin B as a target of SfA in live cells

To begin investigating the targets of SfA by chemical proteomics, we first synthesized two photo-sanglifehrin probes, pSfA1 and pSfA2, by functionalization of SfA at different positions with the minimalist tag, which contains a diazirine for PAL and an alkyne for enrichment (**Figure 1a**).^44^ We designed two pSfA probes to ensure that at least one probe retained activity and to potentially capture a wider range of protein targets (**Extended Data Figure 1a–c**). The effects of the probes on cell viability in Jurkat and K562 cells were assessed as a proxy for immunosuppressive activity in T and B cells, respectively, and had mild immunosuppressive activity, similar to SfA^37^ (**Figure 1b**, **Extended Data Figures 2a–b**). No anti-proliferative activity was observed in A549 cells, indicating that pSfA probes, like SfA,^37^ are not broadly cytotoxic (IC_50_ > 10 µM, **Extended Data Figure 2c**). Using a TR-FRET binding assay, the observed dissociation constants for SfA and the pSfA probes to PPIA and PPIB are generally comparable, although pSfA2 has reduced engagement of PPIA (**Figure 1b**, **Extended Data Figure 3**). Binding of SfA occurs at the highly conserved cyclophilin active site,^45^ which inhibits the enzymatic peptidyl-prolyl cis-trans isomerase (PPI) activity of these enzymes.^46^

**Figure 1:**
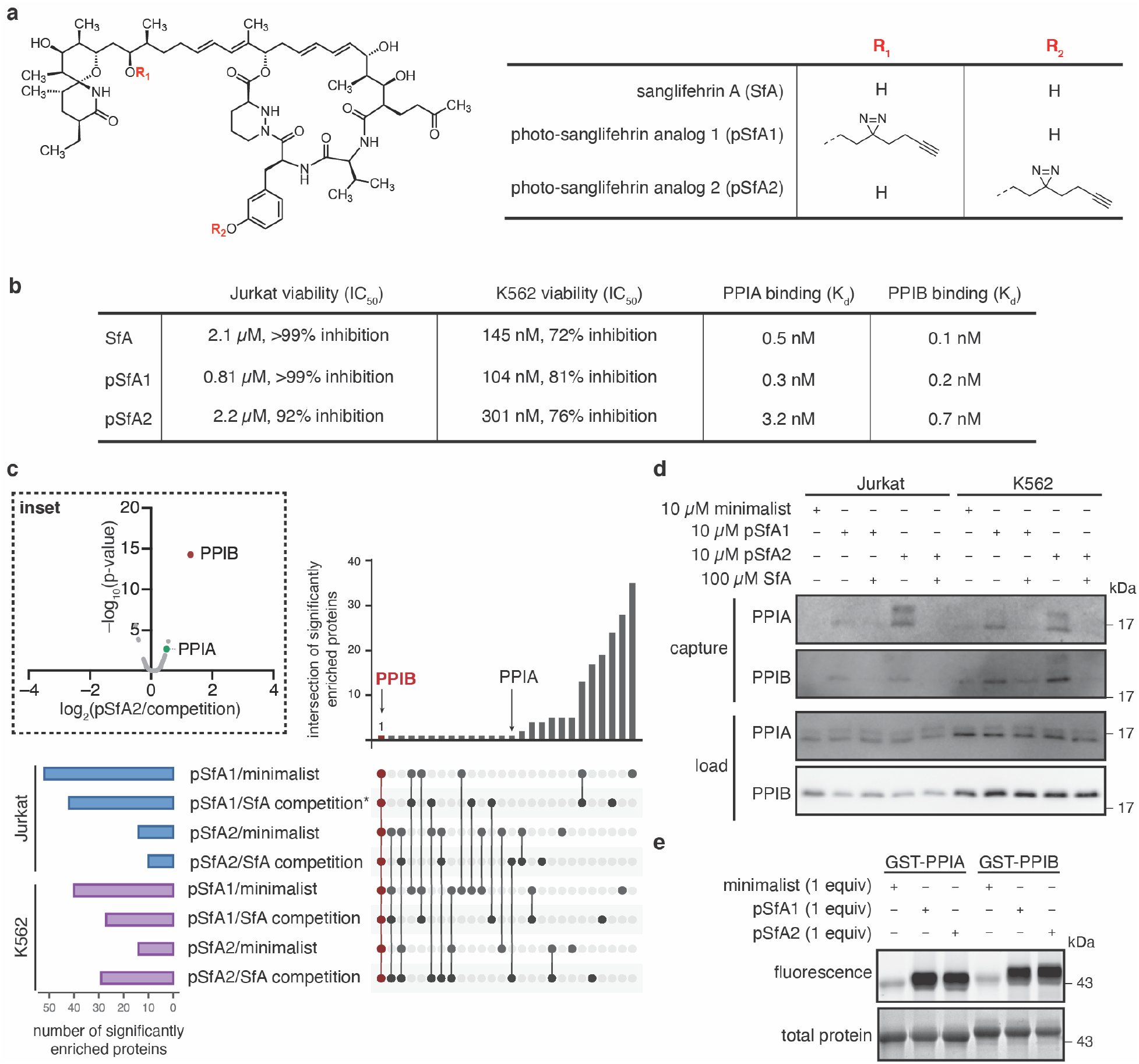
Development of pSfA probes for target identification from live cells. **a,** Structures of SfA, pSfA1, and pSfA2. **b,** Summary of MTT assay results (n = 3) and binding affinities for PPIA and PPIB determined by TR-FRET (n = 2). **c,** Summary of significantly enriched (fold change > 1, p < 0.05) proteins identified by proteomics following treatment with 10 µM pSfA1 or pSfA2, with and without competition with 10x SfA (100 µM), or the minimalist tag alone (10 µM), in Jurkat or K562 cells treated for 30 min prior to photo-affinity labeling (n = 3). * n = 2, due to loss of one sample in the comparison of pSfA1/competition. **Inset**: Example volcano plot showing significant and competitive enrichment of PPIB by pSfA2. **d,** Western blot for PPIA and PPIB after enrichment of pSfA1- or pSfA2-labeled proteins from Jurkat or K562 cells. **e,** In vitro labeling of recombinant GST-PPIA and GST-PPIB with pSfA probes visualized by attachment of Azidefluor488 and in-gel fluorescence.

We next performed chemical proteomics using the pSfA probes in Jurkat and K562 cells in order to identify target proteins of SfA in live cells in an unbiased manner. Proteins labeled and enriched with pSfA1 or pSfA2 from each cell line were considered targets for SfA if they were selectively competed with a ten-fold excess of SfA or were enriched relative to non-specific labeling with the minimalist tag. Among the biological targets of SfA identified by chemical proteomics in live cells (**Table S1)**, surprisingly, only PPIB was enriched significantly [log_2_(fold change) > 1, p < 0.05] across all eight ratios, which represents comparison to SfA competition and background from the minimalist tag alone across two cell lines and two pSfA probes (**Figure 1c, Extended Data Figures 4a–c, Extended Data Figure 5, Table S1**). By contrast, PPIA was significantly enriched by pSfA2 versus competition in Jurkat and K562 cells in two of the eight ratios (**Figure 1c, Extended Data Figures 4a–c, Table S1**). Other PPIases, including mitochondrial PPIF, were observed but not significantly enriched (**Table S1**). The observed cyclophilin interactions were validated by Western blot, which showed that indeed PPIB is labeled to a greater degree than the minimalist tag alone,^47^ and is additionally enriched and competed in both cell lines by pSfA1 and pSfA2 (**Figure 1d**). PPIA is labeled and enriched to a greater extent with pSfA2 after competition with SfA, in alignment with the MS results (**Figure 1c–d**). The pSfA2-treated samples further show a higher molecular weight band for PPIA after enrichment, potentially representing an observable mass shift specifically due to pSfA2 labeling. IMPDH2, a previously identified target of SfA,^37^ showed some labeling by the pSfA probes (**Extended Data Figure 4d**), although it did not meet significance thresholds in MS data (**Table S1**). Despite the preference for PPIB observed in cells, both pSfAs labeled recombinant PPIA and PPIB similarly by in-gel fluorescence (**Figure 1e**). These results indicate that the pSfA probes, and by inference SfA, interact to a greater extent with PPIB than any other cyclophilins in the context of the cellular environment, which is uniquely revealed by the chemical proteomics approach.

### SfA induces secretion of cyclophilin B into the extracellular space

A previous report has described secretion of PPIB upon CsA binding to its catalytic domain.^48^ Since SfA similarly binds to PPIB in live cells, we next sought to assess the effects of SfA on PPIB secretion. Notably, treatment of Jurkat cells with 1 µM SfA produced a decrease in intracellular PPIB protein levels and an associated increase in extracellular PPIB secreted into the conditioned media from these cells within 4 h (**Figure 2a, Extended Data Figure 6a**). No changes in PPIA protein levels in cells or media were observed. CsA treatment had a comparable effect on PPIB secretion under these conditions, SfA-induced PPIB secretion was inhibited by ER transport inhibitors such as brefeldin A, but not by inhibitors of other pathways, such as neddylation (MLN4924), proteasomal degradation (MG132), lysosomal degradation (E64d, pepstatin), or protein synthesis (cycloheximide, **Figure 2a, Extended Data Figure 6b**). These effects were confirmed by fluorescence microscopy for PPIB and the ER marker protein disulfide isomerase (PDI) in HeLa cells, which were used previously to image PPIB in the presence of CsA (**Figure 2b**).^48^ MS analysis of secreted proteins validated the selective increase in extracellular PPIB levels upon SfA or CsA treatment [log2(fold change) > 1, p < 0.05, **Figures 2c, 2d, Table S2**], without affecting cytoplasmic PPIA levels. These data suggest that the secretion of PPIB upon treatment with these cyclophilin binders may result from disruption of native interactions in the ER rather than the formation of a new complex that favors PPIB secretion.

**Figure 2:**
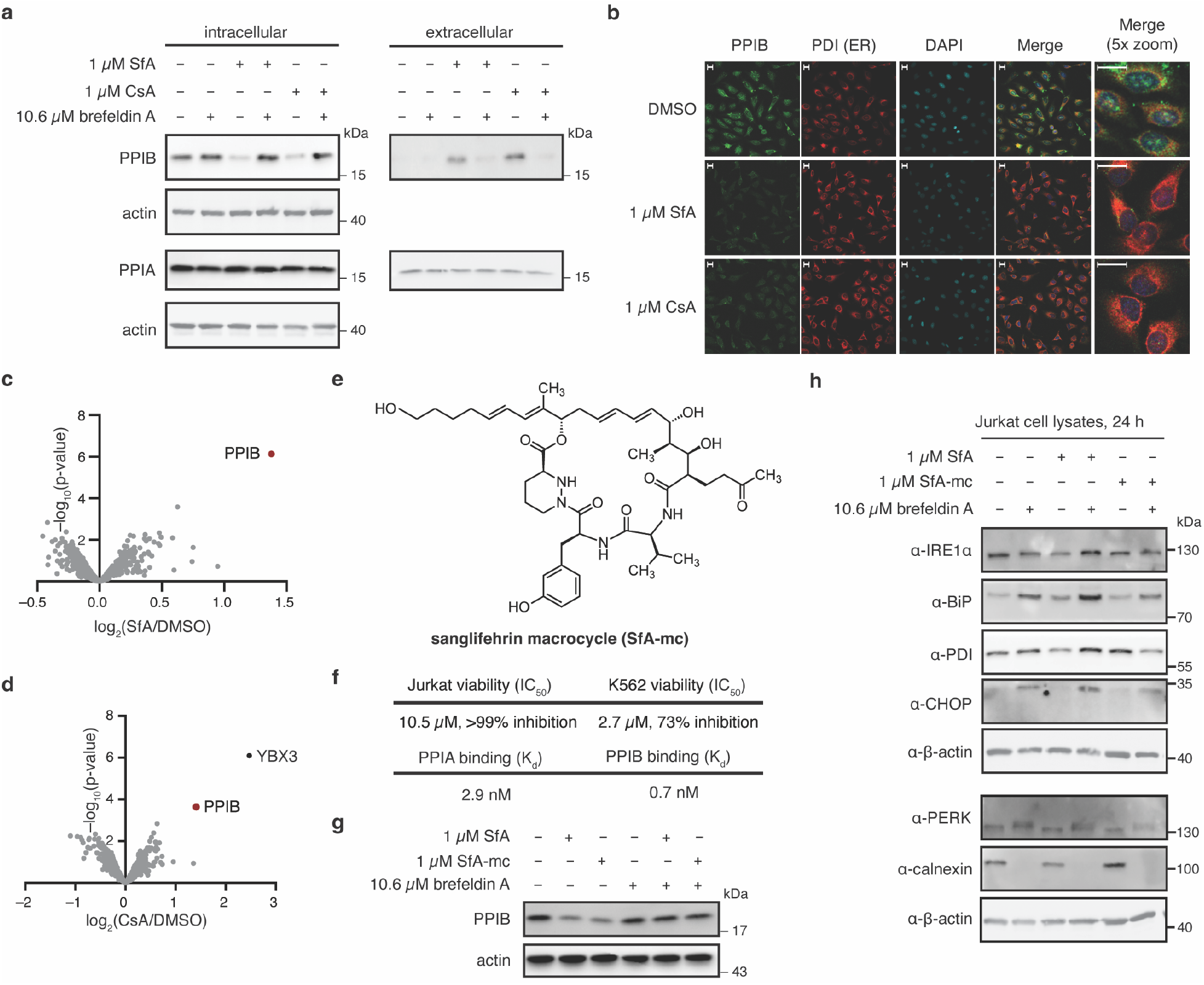
SfA induces secretion of PPIB from cells. **a,** Western blot of intracellular and extracellular PPIB in Jurkat cells treated with 1 µM SfA or 1 µM CsA for 4 h ± 10.6 µM brefeldin A. **b,** Immunofluorescence imaging of PPIB and PDI in HeLa cells treated with 1 µM SfA or 1 µM CsA for 4 h. Scale bars shown (20 µm). **c,** Volcano plot of proteins secreted by Jurkat cells following treatment with 1 µM SfA for 4 h relative to vehicle (n = 3). **d,** Volcano plot of proteins secreted by Jurkat cells following treatment with 1 µM CsA for 4 h relative to vehicle (n = 3). **e,** Structure of sanglifehrin macrocycle (SfA-mc). **f,** Summary of MTT assay results (n = 3) and PPIA and PPIB binding affinities, determined by TR-FRET (n = 2), for SfA-mc. **g,** Western blot of intracellular PPIB in Jurkat cells treated with 1 µM SfA or 1 µM SfA-mc for 4 h ± 10.6 µM brefeldin A. **h**, Western blot of Jurkat cell lysates for ER stress markers after treatment with 1 µM SfA or 1 µM SfA-mc with or without brefeldin A for 24 h (n = 3).

To investigate the role of SfA-induced PPIB secretion in mediating the immunosuppressive effects of SfA, we developed a novel SfA macrocycle functionalized with a primary alcohol (SfA-mc, **Figure 2e**) for comparison to SfA. Prior efforts have found that the SfA macrocycle, obtained by oxidative cleavage of SfA to reveal an α,β-unsaturated aldehyde, retains cyclophilin binding but has reduced immunosuppressive activity.^37,49^ We hypothesized that SfA-mc would provide a useful comparison for determining whether the retention of cyclophilin binding is associated with induction of PPIB secretion but differentiated from immunosuppressive activity. As expected, SfA-mc was less potent in Jurkat and K562 cell viability assays relative to SfA (**Figure 2f, Extended Data Figures 6c–d**). Further, in comparison to SfA, SfA-mc had a similar binding affinity for PPIB but lost some binding affinity for PPIA (**Figure 2f, Extended Data Figure 3**). Excitingly, SfA and SfA-mc induced similar changes in intracellular PPIB levels following treatment of Jurkat cells with 1 µM compound for 4 h (**Figure 2g**). No markers of ER stress were up-regulated in response to SfA or SfA-mc treatment after 24 h, indicating that SfA activity on export of PPIB does not stimulate a broader stress-response signal in cells (**Figure 2h**). By contrast, brefeldin A treatment promoted strong up-regulation of BiP, CHOP, and calnexin markers for ER stress. These data suggest that SfA-mediated PPIB secretion is independent of the immunosuppressive effects exerted by SfA on T cells and B cells and that these effects can be differentiated by utilizing SfA analogs.

### SfA inhibits collagen secretion induced by TGF-β1 in human lung fibroblasts

Given that PPIB is critical for collagen folding,^42,43^ we next focused on evaluating the effect of SfA-induced PPIB secretion in fibrosis, which is characterized by excessive production of collagen type I. PPIB is a major enzyme catalyzing the rate-limiting step of pro-collagen triple helix folding within the collagen prolyl 3-hydroxylation complex, which is comprised of prolyl 3-hydroxylase 1 (P3H1), cartilage-associated protein (CRTAP), and PPIB.^41-43^ We thus reasoned that SfA-mediated PPIB secretion might interfere with collagen type I synthesis by activated myofibroblasts. To investigate this hypothesis, we first studied this mechanism in TGF-β1– activated IMR-90 human lung fibroblasts (**Figure 3a**). TGF-β1 is a potent pro-fibrotic factor that induces fibroblast-to-myofibroblast transdifferentiation during tissue repair and fibrosis, a phenotype characterized by increased synthesis and deposition of collagen type I and other ECM proteins.^50,51^ Unlike quiescent fibroblasts, myofibroblasts express α-smooth muscle actin (αSMA), a protein that confers a hyper-contractile phenotype and allows myofibroblasts to remodel and stiffen the ECM.^1,52^ MS profiling of the secretome of IMR-90 human lung fibroblasts stimulated with TGF-β1 in the presence or absence of SfA showed that SfA treatment reduced protein levels of several collagens in the supernatant of TGF-β1–treated fibroblasts (**Figure 3b, Table S3**). SfA treatment also led to increased PPIB levels in fibroblast supernatants, further validating the induction of PPIB secretion by SfA (**Figures 3b–c, Table S3**).

**Figure 3:**
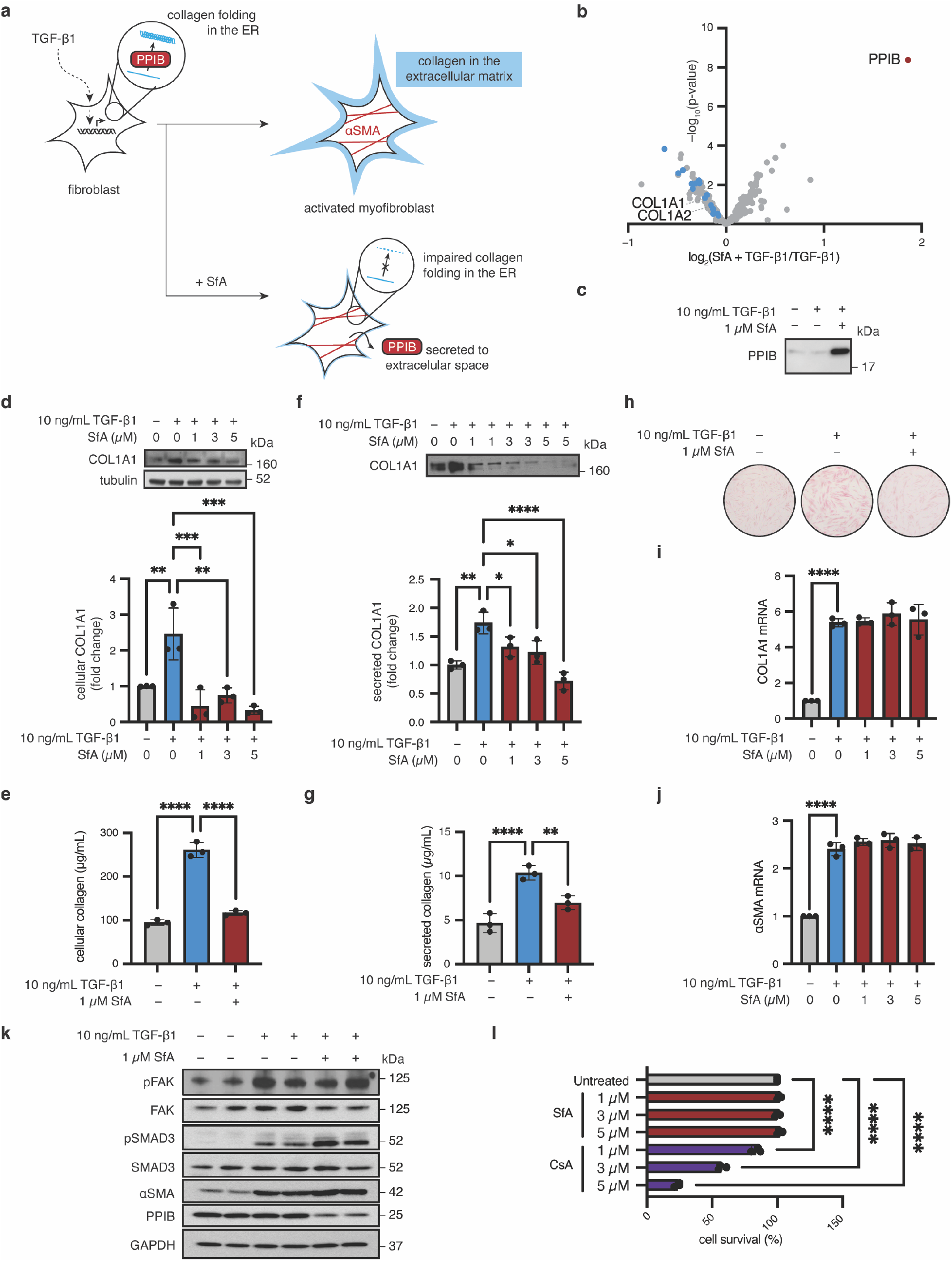
SfA reduces collagen production in an IMR-90 fibroblast model of fibrosis. **a,** Schematic of proposed mechanism of action of SfA in myofibroblasts after TGF-β1–activation. **b,** Secretomics of myofibroblasts treated with SfA. Collagens are highlighted in blue; PPIB is highlighted in red (n = 3). **c,** Western blot for PPIB in conditioned media analyzed in **b**. **d,** Western blot for intracellular collagen (COL1A1) following stimulation with 10 ng/mL TGF-β1 ± 1 µM SfA for 96 h (n = 3). Fold change was calculated by densitometry. **e,** Sircol assay for intracellular collagen following stimulation with 10 ng/mL TGF-β1 ± 1 µM SfA for 96 h (n = 3). **f,** Western blot for extracellular collagen (COL1A1) following stimulation with 10 ng/mL TGF-β1 ± 1 µM SfA for 96 h (n = 3). Fold change was calculated by densitometry. Conditioned media was derived from samples in panel **d**. **g,** Sircol assay measuring extracellular collagen following stimulation with 10 ng/mL TGF-β1 ± 1 µM SfA for 96 h (n = 3). **h,** Intracellular collagen visualized by sircol staining following stimulation with 10 ng/mL TGF-β1 ± 1 µM SfA for 96 h. **i,** Analysis of COL1A1 gene expression following stimulation with 10 ng/mL TGF-β1 ± 1 µM SfA for 96 h (n = 3). **j,** Analysis of αSMA gene expression following stimulation with 10 ng/mL TGF-β1 ± 1 µM SfA for 96 h (n = 3). **k,** Western blot for cellular proteins associated with myofibroblast activation following stimulation with 10 ng/mL TGF-β1 ± 1 µM SfA for 96 h. **l,** Survival as determined by trypan blue staining following treatment with the indicated compounds for 96 h in serum-free media. All graphed data represents means ± standard deviation (SD). Significance was determined by one-way ANOVA followed by pairwise comparisons corrected for multiple comparisons using the Šidák correction. ns = not significant, * = p < 0.05, ** = p < 0.01, *** = p < 0.001, **** = p < 0.0001.

To further investigate whether SfA-induced PPIB secretion regulates collagen type I secretion by fibroblasts, we assessed collagen levels in cell lysates and supernatants by sircol assay and Western blot. As shown in **Figures 3d–h**, SfA treatment significantly reduced both intracellular and extracellular collagen type I (COL1A1) in TGF-β1–treated fibroblasts in a dose-dependent manner. SfA did not affect COL1A1 mRNA levels, consistent with the notion that production of mature, folded collagen type I, but not transcription, is inhibited by SfA (**Figure 3i**). Importantly, SfA not only binds to PPIB and induces its secretion, but also forms a PPIA:SfA binary complex that regulates the cell cycle via inosine-50-monophosphate dehydrogenase 2 (IMPDH2).^37^ To rule out that inhibition of the PPIA:SfA:IMPDH2 pathway by SfA is contributing to the reduction of collagen synthesis in fibroblasts, we tested SfA-mc, which does not form a PPIA:SfA:IMPDH2 complex. SfA-mc potently binds to and induces secretion of PPIB and possesses the same potency as SfA at blocking collagen type I secretion by fibroblasts, further suggesting that SfA-mediated PPIB secretion is the mechanism by which collagen synthesis is reduced (**Extended Data Figure 7a**). In addition, our experiments demonstrated that SfA inhibits collagen synthesis and secretion without affecting αSMA protein or mRNA levels in TGF-β1– activated myofibroblasts, indicating that SfA does not affect the contractility of these cells (**Figures 3j–k**). Further, pro-fibrotic signaling pathways including SMAD2/3 and FAK signaling, which are strongly activated by TGF-β1 in myofibroblasts,^53,54^ are also unaffected by SfA (**Figure 3k**). These results demonstrate the specific activity of SfA, which reduces collagen type I protein levels without affecting upstream pro-fibrotic pathways. Notably, SfA treatment also reduced intracellular PPIB protein levels in TGF-β1–activated myofibroblasts (**Figure 3k**), consistent with our proposed mechanism of action. Moreover, SfA or SfA-mc treatment did not induce myofibroblast apoptosis, in contrast to the toxicity observed in response to CsA treatment (**Figure 3l, Extended Data Figure 7b**). Taken together, our data show that SfA induces PPIB secretion in myofibroblasts, an effect associated with a significant reduction in collagen type I levels *in vitro*. Given the major role of myofibroblasts in the development and progression of fibrotic disease, we next sought to investigate the therapeutic efficacy of SfA at treating fibrosis *in vivo* in a preclinical mouse model.

### SfA reduces established skin fibrosis in the bleomycin mouse model

Our *in vitro* studies showed that SfA inhibits collagen folding and synthesis in myofibroblasts by inducing PPIB secretion, providing a strong rationale for testing the potential anti-fibrotic effects of SfA *in vivo* in a preclinical model. We opted for the well-established bleomycin-induced skin and lung fibrosis mouse model, which is widely used to study the biology of myofibroblasts and to test the efficacy of anti-fibrotic drugs *in vivo*.^55,56^ Using this model, we examined the therapeutic potential of SfA to treat established skin and lung fibrosis when administered therapeutically from day 14 to 28 after the onset of daily bleomycin challenges (**Figure 4a**). Therapeutic administration of SfA (10 mg/kg daily) significantly reduced bleomycin-induced skin fibrosis when compared to vehicle control at day 28 post-bleomycin challenge, as assessed by picrosirius red staining of the skin for collagen, measurement of skin dermal thickness, and skin hydroxyproline content, a biochemical proxy for collagen deposition (**Figures 4b–d**). The bleomycin-induced increase in dermal thickness was reduced by 78% in bleomycin-challenged, SfA-treated mice compared to bleomycin-challenged, vehicle-treated mice. Additionally, the bleomycin-induced increase in skin hydroxyproline was reduced by 58% with SfA treatment, demonstrating potent anti-fibrotic effects of SfA *in vivo*. Immune cells have been also shown to promote fibrosis in this model by stimulating myofibroblast activation via secretion of pro-fibrotic mediators including TGF-β1.^55,57^ To investigate potential anti-inflammatory effects of SfA, we performed flow cytometry analysis of skin tissue from mice treated with or without SfA after saline or bleomycin challenge (**Figure 4e, Extended Data Figure 8a**). Our results demonstrated that SfA treatment significantly reduced the percentage of inflammatory monocytes in the skin of bleomycin-challenged, SfA-treated mice compared to bleomycin-challenged, vehicle-treated mice. There was no significant difference in the percentage of macrophages, CD4+ T cells, or CD8+ T cells that were present in the skin of bleomycin-challenged, SfA-treated mice compared to bleomycin-challenged, vehicle-treated mice (**Figure 4e**). Together, these results suggest that SfA is beneficial in treating established skin fibrosis by reducing collagen levels and inhibiting monocyte infiltration.

**Figure 4:**
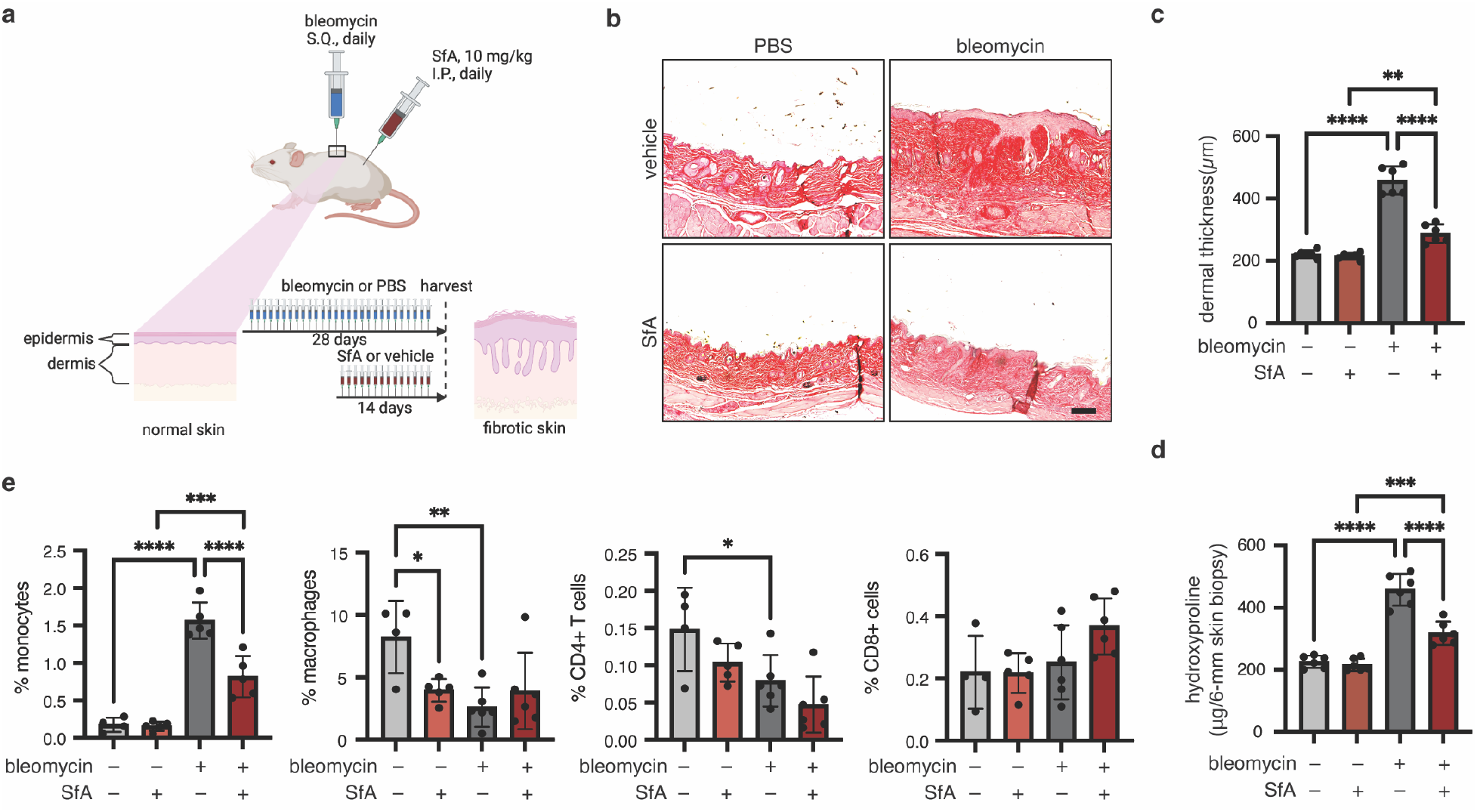
SfA reduces fibrosis and immune activation in a mouse model of bleomycin-induced skin fibrosis. **a,** Schematic of experimental procedure. **b,** Representative images of skin sections stained with picrosirius red to visualize collagen. Scale bar shown (100 µm). **c,** Dermal thickness, as determined by measuring distance between the epidermal–dermal junction and the dermal–fat junction (n = 6). **d,** Collagen content in skin-punch samples, as determined by hydroxyproline assay (n = 6). **e,** Characterization of immune cells in skin biopsy samples. All graphed data represents means ± SD (n = 4–6). **c** and **d** were analyzed using a two-way ANOVA followed by pairwise comparisons corrected for multiple comparisons using the Turkey correction. **e** was analyzed using one-way ANOVA followed by pairwise comparisons corrected for multiple comparisons using the Šidák correction. ns = not significant, * = p < 0.05, ** = p < 0.01, *** = p < 0.001, **** = p < 0.0001.

### SfA reduces established lung fibrosis in the bleomycin mouse model

As subcutaneous injection of bleomycin leads to concomitant pulmonary fibrosis in mice (**Figure 5a**),^58^ we next examined the lungs of mice treated with or without therapeutic SfA at the same timepoints used for the study of skin fibrosis. Therapeutic administration of SfA significantly reduced histological measures of fibrosis in the lungs when compared to vehicle control at day 28 post-bleomycin challenge, as assessed by picrosirius red staining (**Figure 5b**). In addition, hydroxyproline levels in lung tissue, indicative of collagen type I content, were reduced in bleomycin-challenged, SfA-treated mice compared to bleomycin-challenged, vehicle-treated mice (**Figure 5c**). Notably, SfA treatment also reduced the increased alveolar–capillary barrier permeability induced post-bleomycin challenge, as determined by reduction in total protein levels in the bronchoalveolar lavage (BAL) fluid (**Figure 5d**). In addition, SfA treatment also reduced the number of inflammatory monocytes and macrophages in the BAL induced by bleomycin injury, further validating the anti-inflammatory effects of SfA (**Figure 5e, Extended Data Figure 8b**). Further, SfA did not affect the percentage of CD4+ T cells although it did increase the number of CD8+ T cells in fibrotic lungs. Consistent with our proposed mechanism of action SfA, treatment resulted in near-complete loss of PPIB from lung tissue with no changes in PPIA levels (**Figure 5f**),. These results indicate that SfA treats established lung fibrosis by reducing collagen levels and inhibiting monocyte infiltration; SfA treatment was also associated with reduced tissue PPIB levels. To further assess our proposed mechanism of action *in vivo*, we next generated fibrotic precision-cut lung slices (PCLSs) from transgenic collagen-GFP reporter mice (Col-GFP) subjected to our bleomycin lung fibrosis model (**Figure 5g**). In this 3D ex vivo model of lung fibrosis, thin slices freshly prepared from fibrotic tissues represent a powerful tool to study fibrosis mechanism and testing anti-fibrotic responses to novel drug compounds ^59,60^. Our results indicate that treatment of murine fibrotic PCLS with SfA for 2 days reduced collagen type I levels in GFP+ fibrotic fibroblasts directly isolated by FACS from PCLSs (**Figure 5g, 5h**). Of note, COL1A1 mRNA levels were not modulated by SfA treatment (**Figure 5i**), further indicating that SfA prevents collagen synthesis without affecting collagen mRNA transcription. In addition, SfA treatment resulted in reduced soluble collagen type I secretion from PCLS, as assessed in PCLS-derived supernatant (**Figure 5j**). Taken together, our data support PPIB, as engaged by SfA, as a novel therapeutic target for the treatment of skin and lung fibrosis.

**Figure 5:**
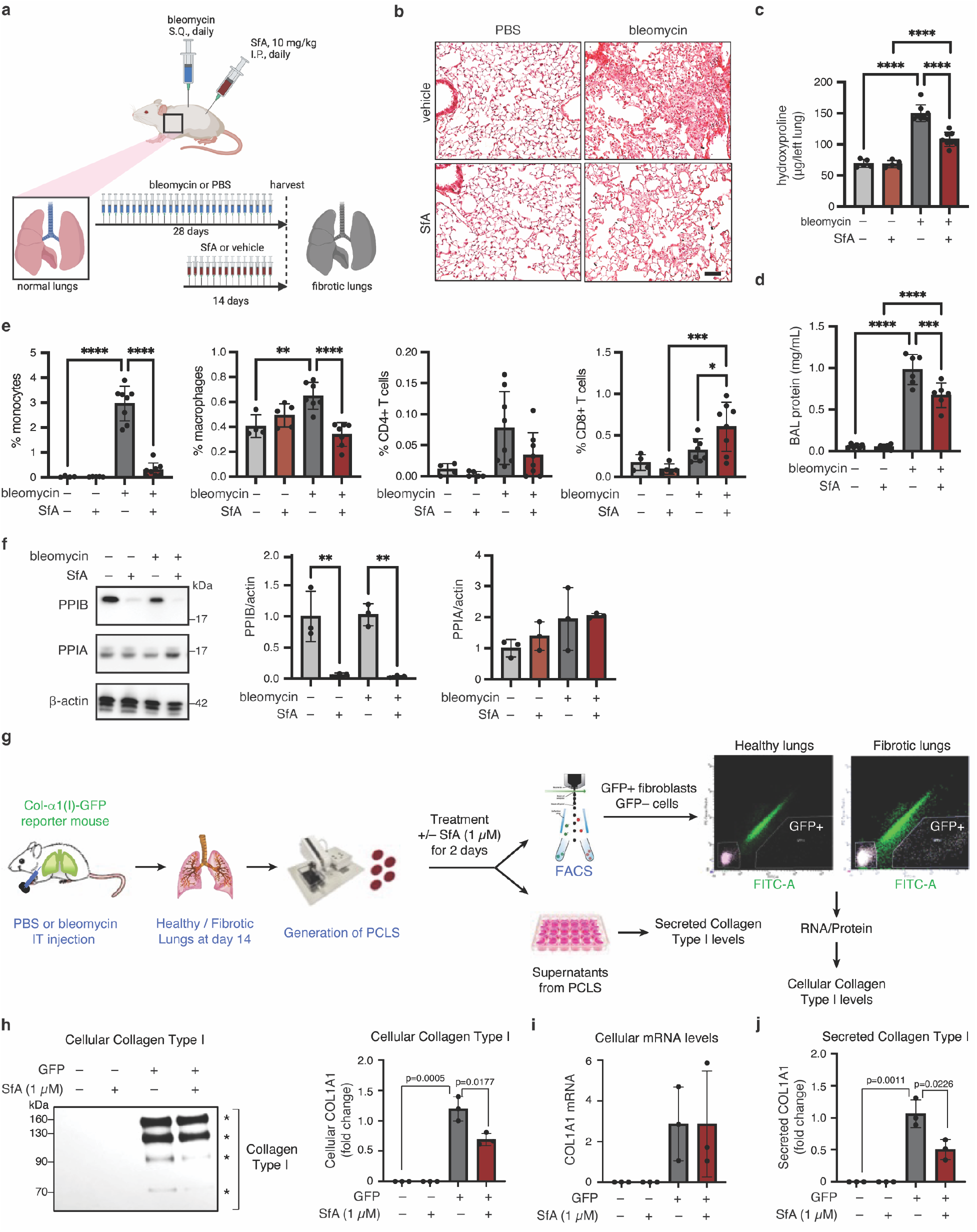
SfA reduces fibrosis and immune activation in a mouse model of bleomycin-induced lung fibrosis. **a,** Schematic of experimental procedure. **b,** Representative images of lung sections stained with picrosirius red to visualize collagen. Scale bar shown (100 µm). **c,** Collagen content in left lungs, as determined by hydroxyproline assay (n = 5–8). **d,** Vascular leak assay, as determined by BCA assay for protein content in bronchioalveolar lavage (BAL) supernatant (n = 6). **e,** Characterization of immune cells in BAL (n = 4–8). **f,** Western blot for PPIA and PPIB in lung cell lysates (n = 3). **g.** Generation of fibrotic precision cut lung slices (PCLS) from transgenic collagen-GFP reporter mice (Col-GFP) at day 14 post-bleomycin challenge. PCLS were treated with or without SfA (1 µm) for 2 days (n = 3). Collagen type I protein and mRNA levels were assessed in GFP– cells and GFP+ fibroblasts sorted by FACS from PCLS by Western blot **(h)** and real time PCR **(i)**, respectively (n = 3). Secreted collagen type I was assessed in PCLS supernatants by sircol assay **(j)** (n = 3). All graphed data represents means ± SD. **c**, **d**, **f, h, i and j** were analyzed using one-way ANOVA followed by pairwise comparisons corrected for multiple comparisons using the Šidák correction. **e** was analyzed using a two-way ANOVA followed by pairwise comparisons corrected for multiple comparisons using the Turkey correction. ns = not significant, * = p < 0.05, ** = p < 0.01, *** = p < 0.001, **** = p < 0.0001.

### SfA inhibits collagen type I secretion by precision-cut lung slices and fibrotic lung fibroblasts from lung tissue of patients with IPF

To determine the relevance of this mechanism in human disease, we assessed collagen type I secretion from PCLSs prepared from explanted lung tissue from patients with IPF (n = 4 individual patients; n ≥ 3 slices per patient/treatment) treated with SfA (1 μM) or vehicle for 3 days (**Figure 6a–c**). Consistent with results observed *in vitro* and in the preclinical mouse model, SfA treatment significantly reduced collagen type I protein secretion compared to vehicle control, an effect associated with PPIB secretion (**Figure 6d–g**). To further investigate anti-fibrotic effects of SfA in IPF, we next assessed the effects of SfA on primary human fibroblasts isolated from IPF patients (n = 3) and healthy controls (n = 3). The fibrotic fibroblasts isolated from IPF patients secreted slightly higher levels of collagen type I into culture medium compared to normal lung fibroblasts *in vitro* using our cultured conditions (**Figure 6h**). Treatment with SfA (1 µM) significantly reduced collagen type I levels in the supernatant of IPF fibroblasts, while treatment of healthy control fibroblasts with SfA did not have a significant effect on collagen type I levels (**Figure 6h**). Consistent with our *in vitro* studies in IMR-90 cells, SfA (1 µM) treatment did not modulate IPF fibroblast contractility, assessed by αSMA protein levels, or TGF-β1/SMAD signaling (**Figure 6i**). Taken together, our human studies confirmed our proposed mechanism by which SfA-induced PPIB secretion inhibits collagen type I secretion from IPF fibroblasts without affecting other fibroblast functions or fibrogenic TGF-β1 signaling.

**Figure 6:**
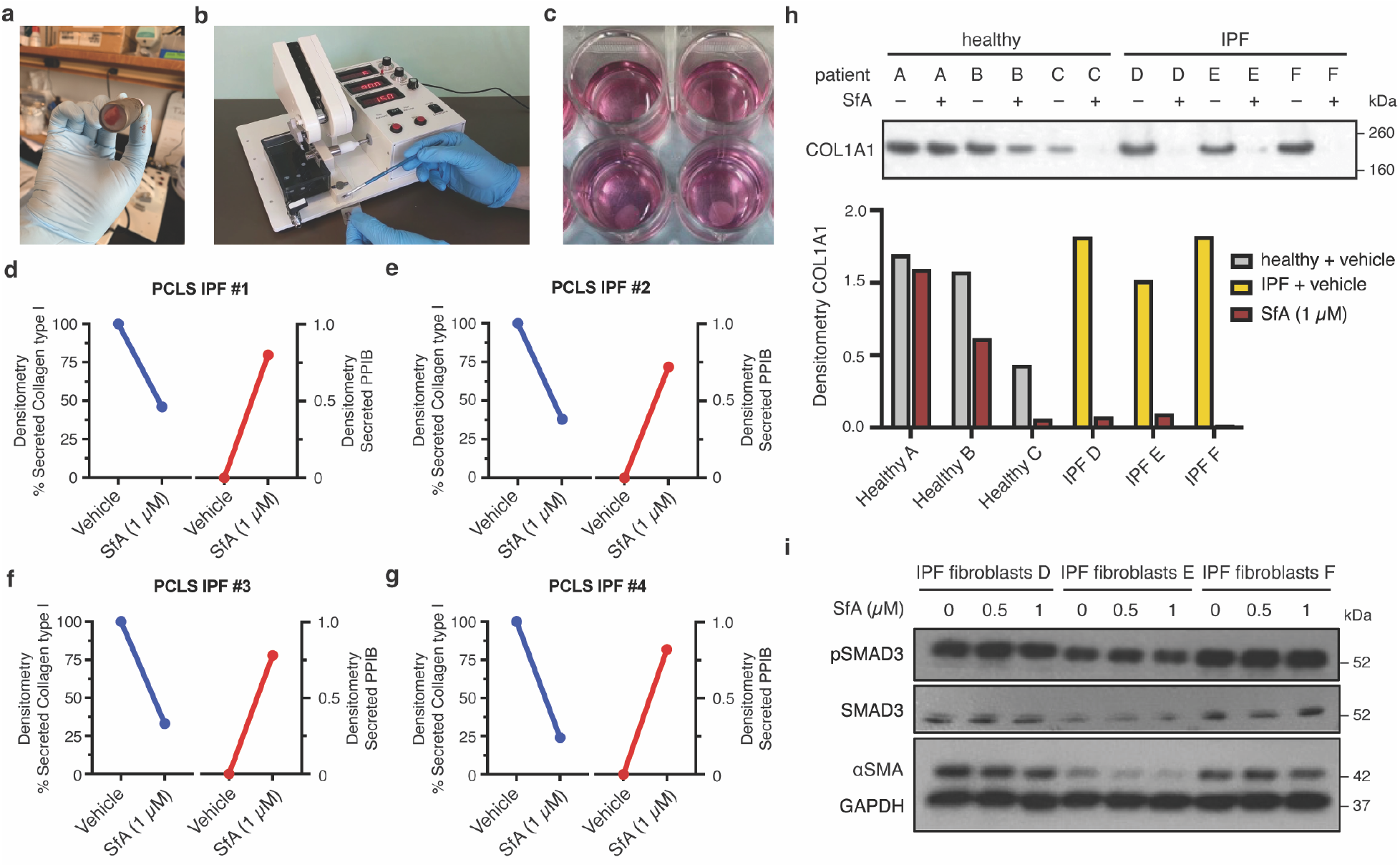
SfA reduces collagen secretion from primary fibrotic fibroblasts. **a,** Generation of precision cut lung slices (PCLS) from explanted lung tissue isolated from patients with idiopathic pulmonary fibrosis (IPF). Lung tissue was obtained in 1 cm blocks (**a**), then sliced (200–300 µm thick) using a Compresstome® VF-310-0Z (**b**). PCLSs were then treated with or without SfA (1 µM) for 72 h in culture (n = 4) (**c**). **(d–g)** Analysis of collagen and PPIB secretion from PCLSs prepared from IPF patient lung tissue ± 1 µM SfA in supernatants (n = 4). **h,** Western blot for collagen type I secreted by primary human fibroblasts isolated from healthy controls and IPF patients ± 1 µM SfA over 96 h in cell supernatants. **i,** Western blot for α-smooth muscle actin (αSMA) and phosphoSMAD3/SMAD3 signaling in IPF fibroblasts ± SfA over 96 h in cell lysates. GAPDH was used as loading control. **d–g** was analyzed with a one-sample t-test against a hypothetical value of 1.

## Discussion

Skin and lung fibrosis are lethal components and hallmarks of human fibrotic diseases such as SSc and IPF.^1,5,7,8^ The scar tissue that builds up in fibrotic disease is primarily composed of collagen type I, which is synthesized and secreted by myofibroblasts activated by pro-fibrotic mediators such as TGF-β1.^1^ Inhibition of TGF-β1–induced collagen type I transcription by myofibroblasts is thought to be the major mechanism targeted by pirfenidone and nintedanib, the only FDA-approved anti-fibrotic agents. However, the anti-fibrotic effects of these drugs are modest and associated with low tolerability due to off-target side effects. The development of novel anti-fibrotic agents with greater selectivity and improved efficacy requires identification of new drug targets and elucidation of novel mechanism(s) of action. Here, we applied PAL and chemical proteomics to unravel a novel mechanism of action for SfA in fibrosis. Our results demonstrate that SfA binds to and induces secretion of PPIB, an ER-resident chaperone involved in collagen type I folding and maturation. Our studies in Jurkat and K562 cells revealed that PPIB is the major target of SfA in live cells, which induces the secretion of PPIB into the extracellular space and depletes intracellular PPIB. Depletion of PPIB is reminiscent of a PPIB knockdown. Knockdown of proteins is an emerging therapeutic modality that has been achieved by preventing protein expression with siRNA or promoting protein degradation through targeted protein degradation and is achieved through a third mechanism of secretion here. Comparison of SfA and SfA-mc, an analog with reduced immunosuppressive effects that similarly induces PPIB secretion, indicates that PPIB secretion can be separated from the immunosuppressive effects of SfA in Jurkat or K562 cells.

Given the importance of PPIB in collagen type I folding and maturation, we hypothesized that SfA might affect collagen maturation in fibroblasts. Consistent with our proposed mechanism, we showed that SfA treatment reduces collagen type I levels in TGF-β1–activated myofibroblasts by promoting secretion of PPIB into the extracellular space. These results translated to a mouse model of bleomycin-induced skin and lung fibrosis, where therapeutic SfA treatment (10 mg/kg daily) resulted in reduced fibrosis as determined by histological and biochemical measures. Importantly, PPIB levels in lung tissue were drastically reduced with SfA treatment, supporting *in vivo* translation of the mechanism observed *in vitro.* SfA additionally dampened the innate immune response as indicated by reduced monocyte infiltration, suggesting a dual mechanism acting both on collagen maturation, through induction of PPIB secretion, and on the innate immune response by yet unknown mechanisms. Further, in PCLSs and fibroblasts isolated from patients with IPF, SfA treatment reduced collagen secretion, suggesting that the effects observed in the mouse model are translatable to IPF, a human disease sorely in need of additional treatments.

From a translational perspective an important future direction is to evaluate whether compounds without immunosuppressive activity, such as nonimmunosuppressive SfA and CsA analogs,^61^ are preferable for anti-fibrotic applications. Our data shows that SfA has both anti-inflammatory and anti-fibrotic effects of SfA in a preclinical mouse model, which may be separable as shown with SfA-mc. Therapeutic targeting of cyclophilins by non-immunosuppressive analogs derived from SfA and CsA, previously developed as anti-virals,^61^ has been shown to have anti-fibrotic effects in the CCl_4_ model of liver fibrosis and in a mouse model of nonalcoholic steatohepatitis (NASH).^62,63^ In this regard, a novel non-immunosuppressive pan-cyclophilin inhibitor CRV431 (Rencofilstat), a CsA analog that is described as pan PPIA, PPIB, PPIF, and PPIG inhibitor, has been recently entered into clinical development. Hepion Pharmaceuticals, Inc has recently shown that rencofilstat is safe in a Phase I human trial and is planning to test its anti-fibrotic effects in a Phase 2B trial in NASH (ASCEND: HEPA-CRV431-207). SfA shows promising activity *in vivo* and further structure-activity relationship studies in addition to SfA-mc may yield optimized molecules.

Additionally, SfA is a pan-cyclophilin inhibitor and therefore a role for other PPIases in the overall mechanism of action of SfA cannot be completely excluded without the development of more selective SfA analogs. Nonetheless, our studies demonstrate the role of PPIB in regulating collagen type I synthesis by myofibroblasts during the development of lung and skin fibrosis *in vivo*. This mechanism is partly supported by genetic studies demonstrating reduced collagen type I crosslinking in PPIB-deficient mice, which also develop features of osteogenesis imperfecta.^42,43,64,65^ Notably, though excessive TGF-β1 signaling is a common mechanism in these mouse models of osteogenesis imperfecta, these mice do not develop organ fibrosis,^66^ which is consistent with the notion that PPIB may be required for TGF-β1 signaling to initiate the development of fibrosis. By contrast, studies with the SfA-derived pan-cyclophilin inhibitor GS-642362, which targets PPIA, PPIB, and PPIF, in the unilateral ureteric obstruction (UUO) mouse model showed inhibition of renal fibrosis by preventing tubular epithelial cell death and neutrophil infiltration.^67^ PPIB-deficient mice are viable and partially protected from inflammation in the UUO model at day 7, although fibrosis was not assessed at later time points,^68^ while efforts with PPIA-deficient mice show reduced inflammation in the bilateral renal ischemia/reperfusion injury (IRI) model but are not protected from renal fibrosis in the UUO model, suggesting that PPIA regulates inflammation, but not fibrosis.^69^ Furthermore, PPIF-deficient mice showed protection from renal fibrosis in the UUO model due to reduction in tubular epithelial cell apoptosis^70^ and TGF-β1– induced collagen type I expression in fibroblasts isolated from these mice was comparable to that observed in wild-type fibroblasts, suggesting PPIF does not play a direct role in fibrogenic responses. Further studies aided by the development of more selective PPIB inhibitors are needed to elucidate the molecular underpinnings linking PPIB, TGF-β1, and organ fibrosis *in vivo*. A strategy to generate isoform-selective cyclophilin inhibitors was recently demonstrated by engineering macrocycle scaffolds.^71^ Our studies indicate that PPIB is a novel anti-fibrotic target and motivates future optimization of compounds to selectively target PPIB in collagen type I production.

In summary, our work shows that the immunosuppressive natural product SfA exerts anti-fibrotic effects by depleting intracellular PPIB, interfering with collagen maturation and reversing bleomycin-induced skin and lung fibrosis in a mouse model. This mechanism represents a novel strategy for treating organ fibrosis by interfering with collagen type I maturation in myofibroblasts. We uncovered this mechanism through profiling of SfA targets by PAL and chemical proteomics, which demonstrated that PPIB is the major cellular target of SfA and revealed that SfA induces PPIB secretion. Incorporation of drugs acting through this novel mechanism into the currently paltry arsenal of anti-fibrotic drugs has the potential to usher in an era of improved outcomes for patients suffering from fibrosis.

## Data availability

Proteomics data have been deposited in the PRIDE repository under identifiers PXD029540, PXD029541, and PXD031010.

## Author contributions

H.A.F., M.C., H.H., F.K., and R.T.L. performed experiments and analyzed data. C.C. synthesized compounds. K.E.B. provided samples of primary tissue. C.M.W. and D.L. supervised the study. H.A.F., C.M.W., and D.L. wrote the manuscript with input from other authors.

## Acknowledgements

We thank S. Trager, B. Budnik, and R. Robinson from the Harvard University Mass Spectrometry and Proteomics Resource Laboratory for technical support. Sanglifehrin A was a generous gift from Novartis.

## Figures

**Extended Data Figure 1:**
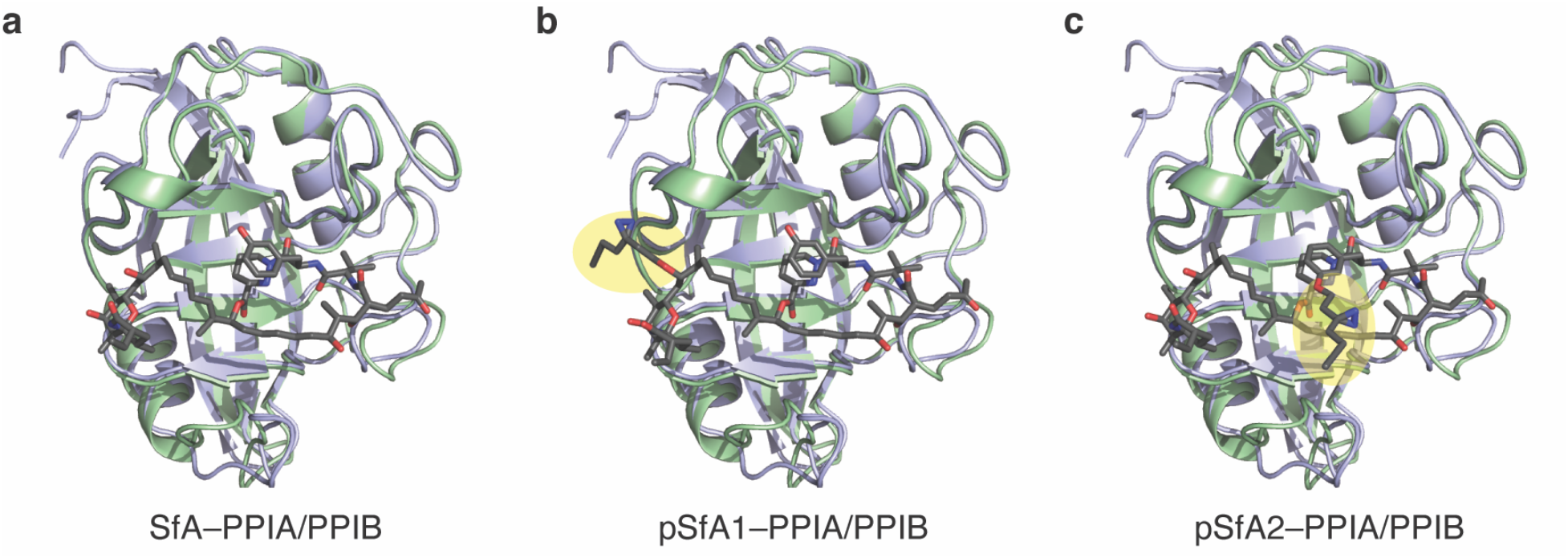
Model of SfA and analogs pSFA1 or pSFA2 overlaid on PPIA and PPIB. **a,** Crystal structure of the SfA–PPIA complex (green, PDB: 1YND) aligned with PPIB (blue, PDB: 3ICI). **b**, Model of pSfA1 overlaid on PPIA and PPIB. **c**, Model of pSfA2 overlaid on PPIA (green) and PPIB (blue). Photo-affinity tag highlighted in yellow. Models of pSfA1 and pSfA2 were derived from SfA and the chemical modification was minimized in Molecular Operating Environment 2020.

**Extended Data Figure 2:**
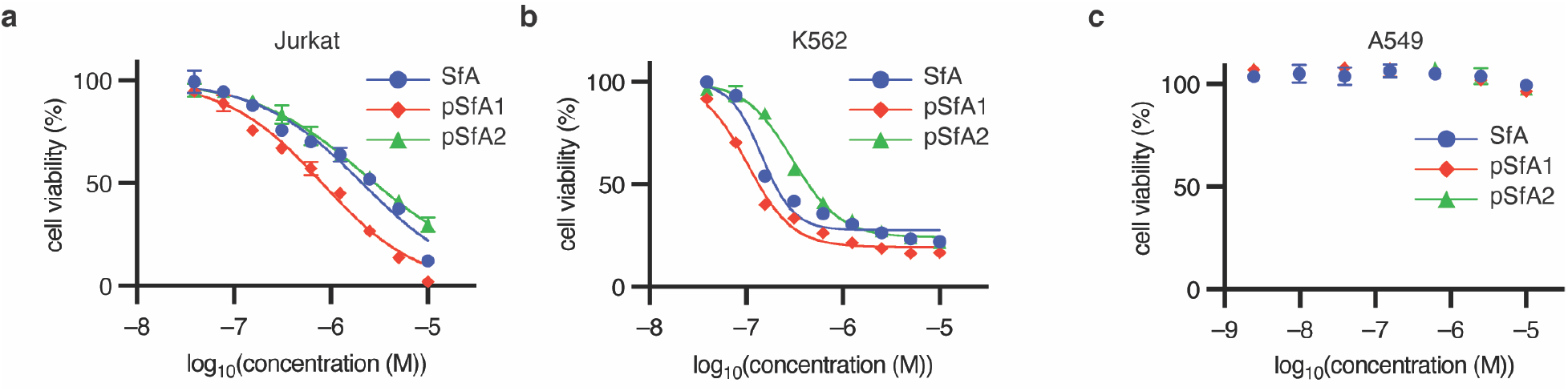
Effect of indicated compounds on Jurkat (**a**), K562 (**b**), and A549 (**c**) cell viability, as determined in a 72 h MTT assay (n = 3). MTT data is shown as means ± SD, with 3 replicates per condition.

**Extended Data Figure 3:**
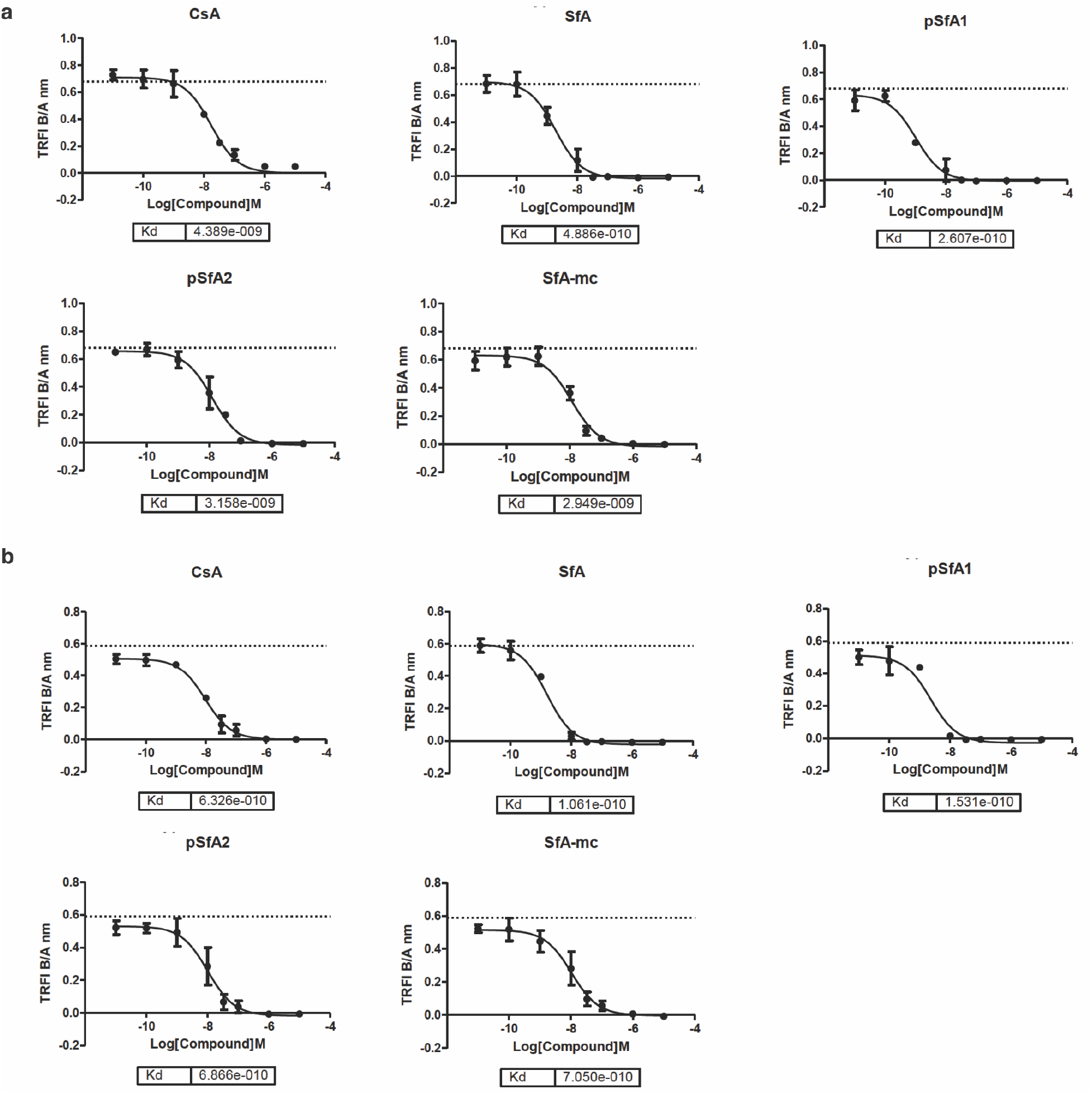
Determination of dissociation constants for SfA, pSfA1, pSfA2, and SfA-mc to PPIA (**a**) and PPIB (**b**), using a standard protocol performed by Eurofins (n = 2). Dissociation constants were determined relative to CsA in a competitive TR-FRET assay, with each compound assayed over a concentration range of 0.01 nM–10 µM.

**Extended Data Figure 4:**
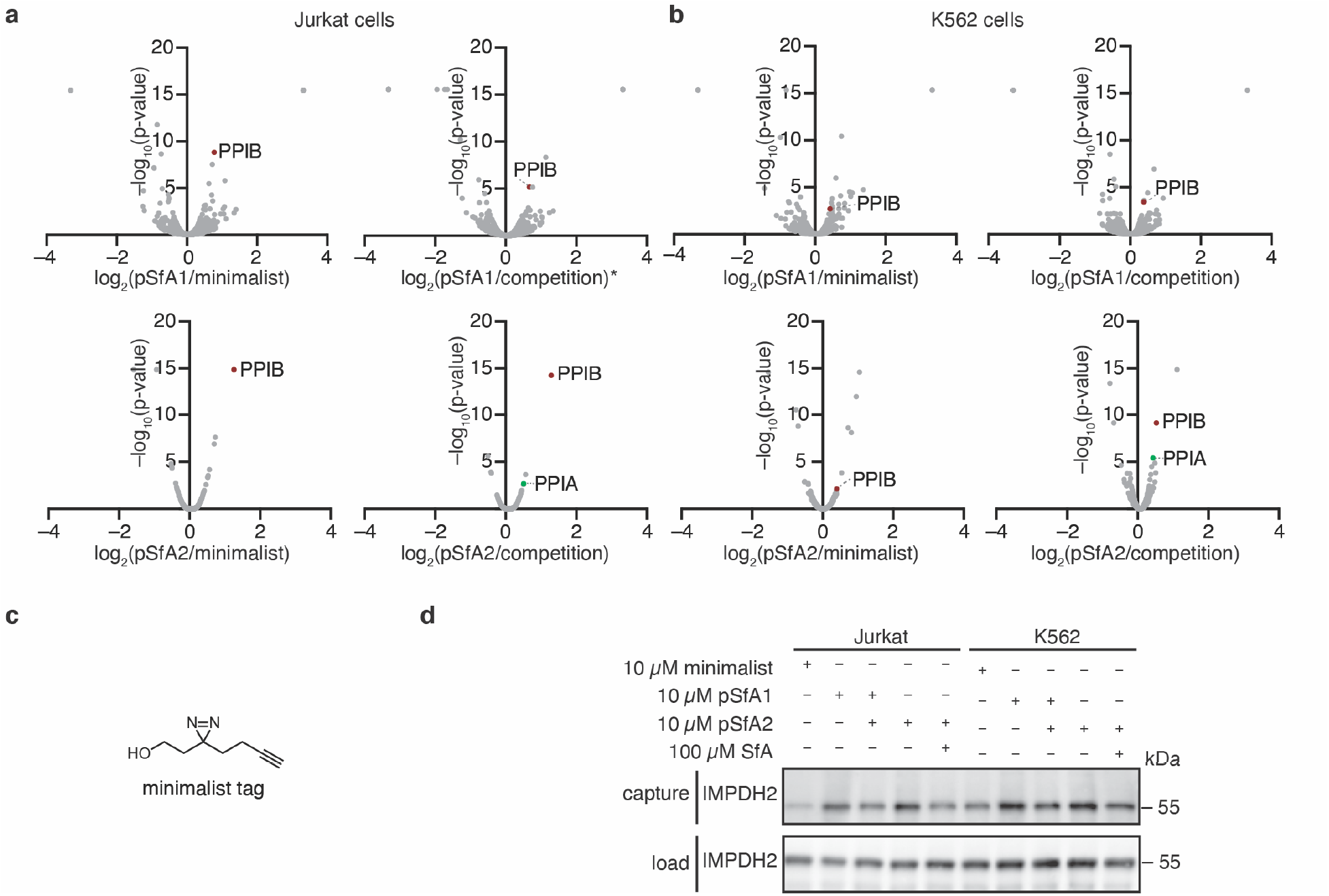
Photo-affinity labeling with sanglifehrin analogs and chemical proteomics analysis. **a,** Volcano plots of chemoproteomics data from Jurkat cells (n = 3). PPIB is marked in red on each plot. PPIA is marked in green in ratios where it was significantly enriched. * n = 2, due to loss of one sample in the comparison of pSfA1/competition. **b,** Volcano plots of chemoproteomics data from K562 cells (n = 3). PPIB is marked in red on each plot. PPIA is marked in green in ratios where it was significantly enriched. **c,** Structure of the minimalist tag used as a negative control in chemoproteomics experiments. **d,** Western blot of enrichment of IMPDH2 with pSfA1 and pSfA2 from Jurkat and K562 cells.

**Extended Data Figure 5:**
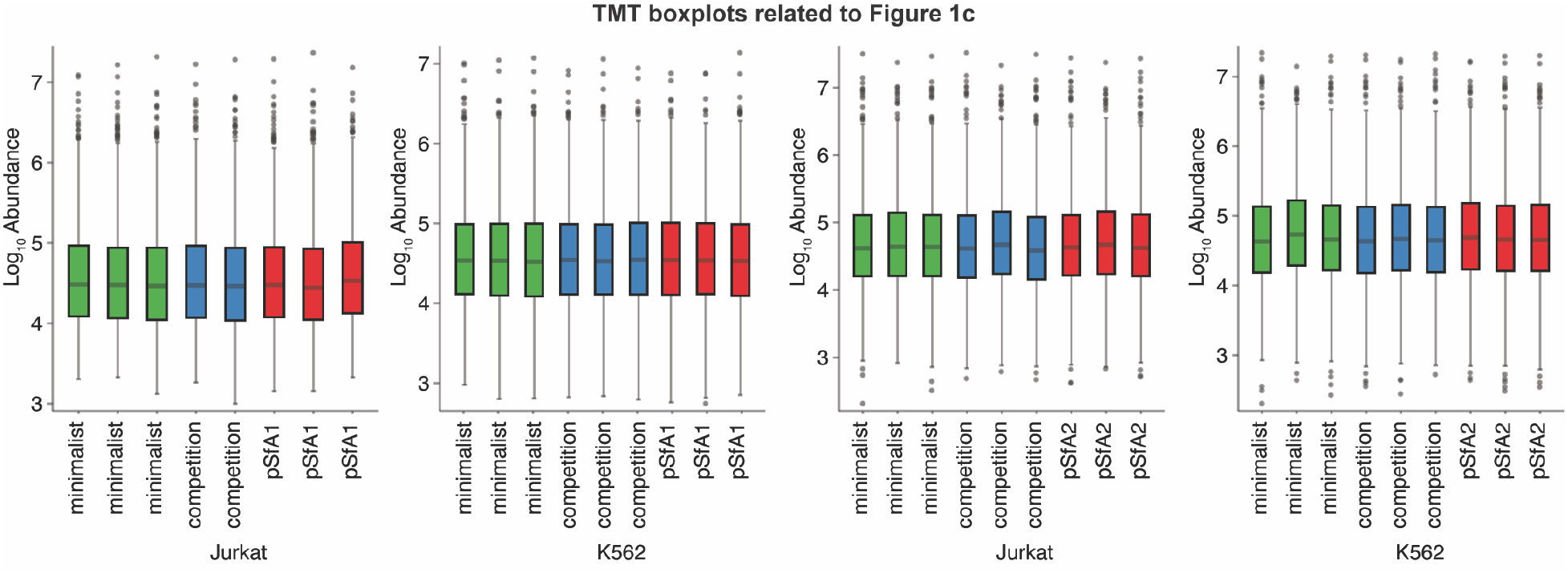
Boxplot of protein level normalization using TMT. Normalization was performed against the total peptide amount in Proteome Discoverer 2.4.

**Extended Data Figure 6:**
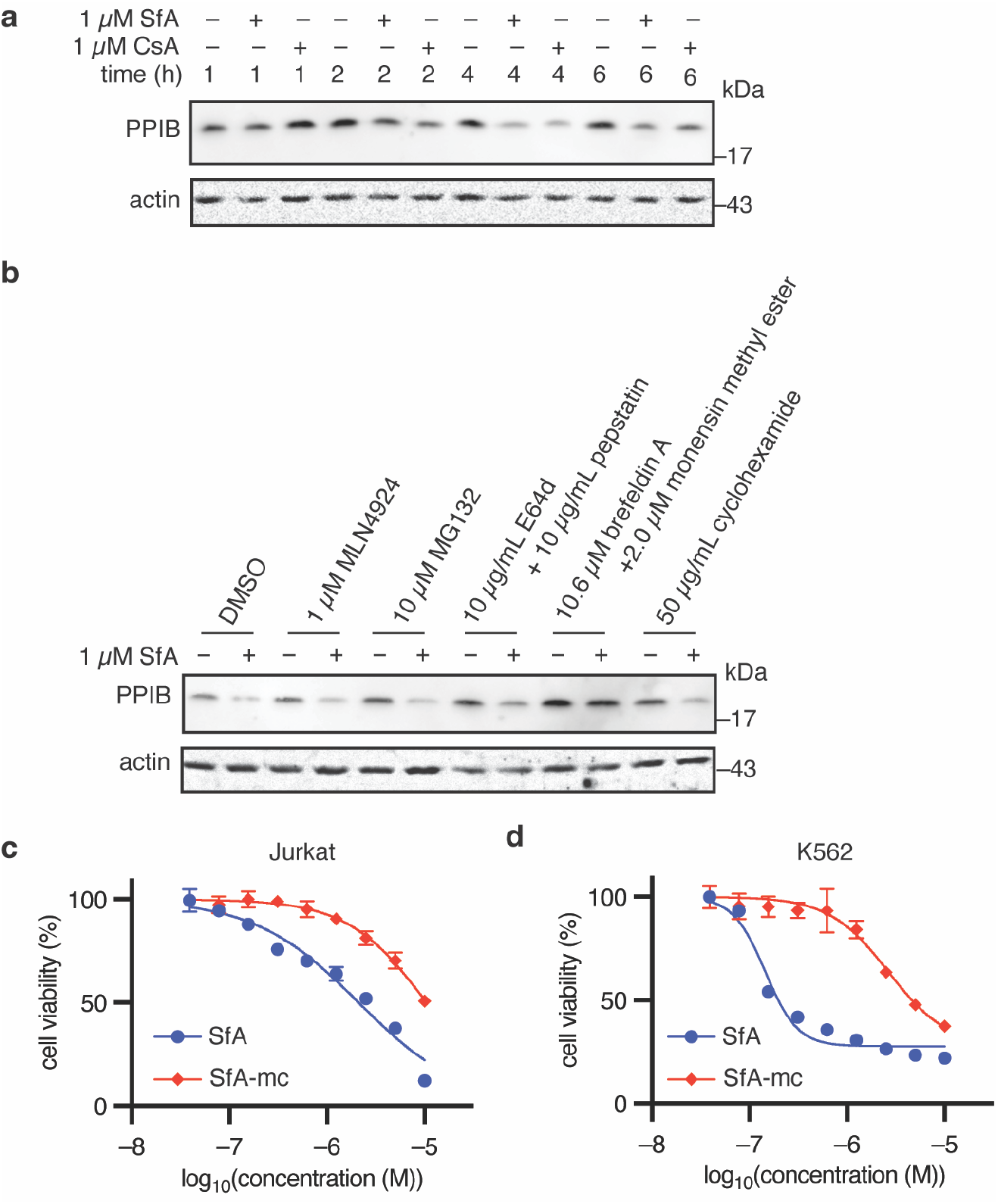
Comparison of SfA, CsA, and SfA-mc on PPIB levels and cell viability. **a,** Western blot of intracellular PPIB in Jurkat cells treated with 1 µM SfA or 1 µM CsA for the indicated time. **b,** Western blot of intracellular PPIB in Jurkat cells pretreated with the indicated inhibitors for 1 h prior to treatment with 1 µM SfA for 4 h. **c,** Activity of SfA and SfA-mc in a 72 h MTT assay in Jurkat cells. SfA data is replicated from Extended Data Figure 2a (n = 3). **d,** Activity of SfA and SfA-mc in a 72 h MTT assay in K562 cells (n = 3). SfA data is replicated from Extended Data Figure 2b (n = 3). MTT data is shown as means ± SD, with 3 replicates per condition.

**Extended Data Figure 7:**
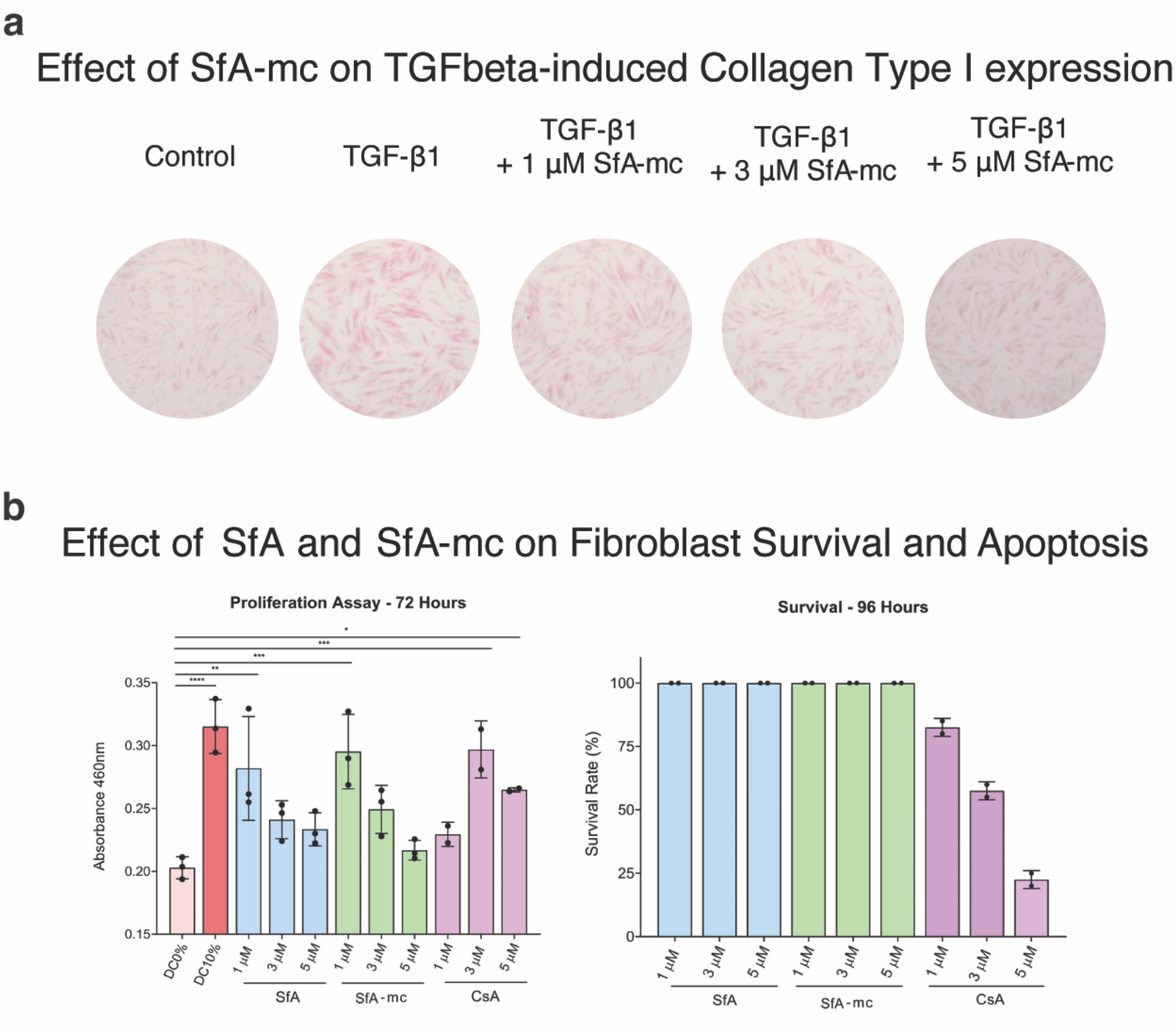
SfA-mc reduces collagen production in an IMR-90 fibroblast model of fibrosis. **a,** Intracellular collagen visualized by sircol staining following stimulation with 10 ng/mL TGF-β1 ± 1 µM SfA-mc for 96 h. **b,** Survival as determined by trypan blue staining following treatment with the indicated compounds for 96 h in serum-free media. All graphed data represents means ± standard deviation (SD). Significance was determined by one-way ANOVA followed by pairwise comparisons corrected for multiple comparisons using the Šidák correction. ns = not significant, * = p < 0.05, ** = p < 0.01, *** = p < 0.001, **** = p < 0.0001.

**Extended Data Figure 8:**
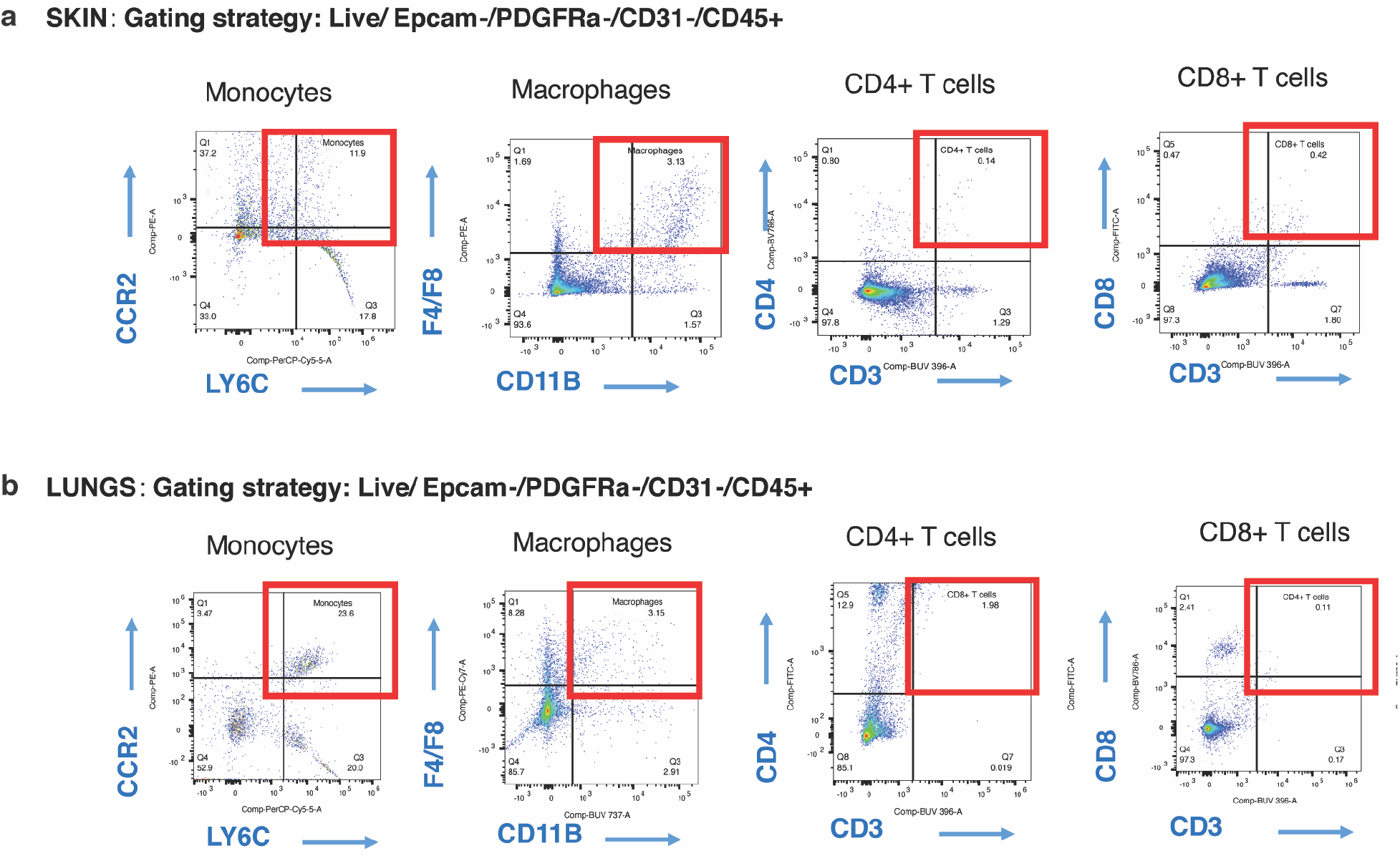
Gating schemes for flow cytometry data. Gating schemes for data shown in Fig. 4e (**a**) and 5e (**b**).

**Supplementary Figure 1:**
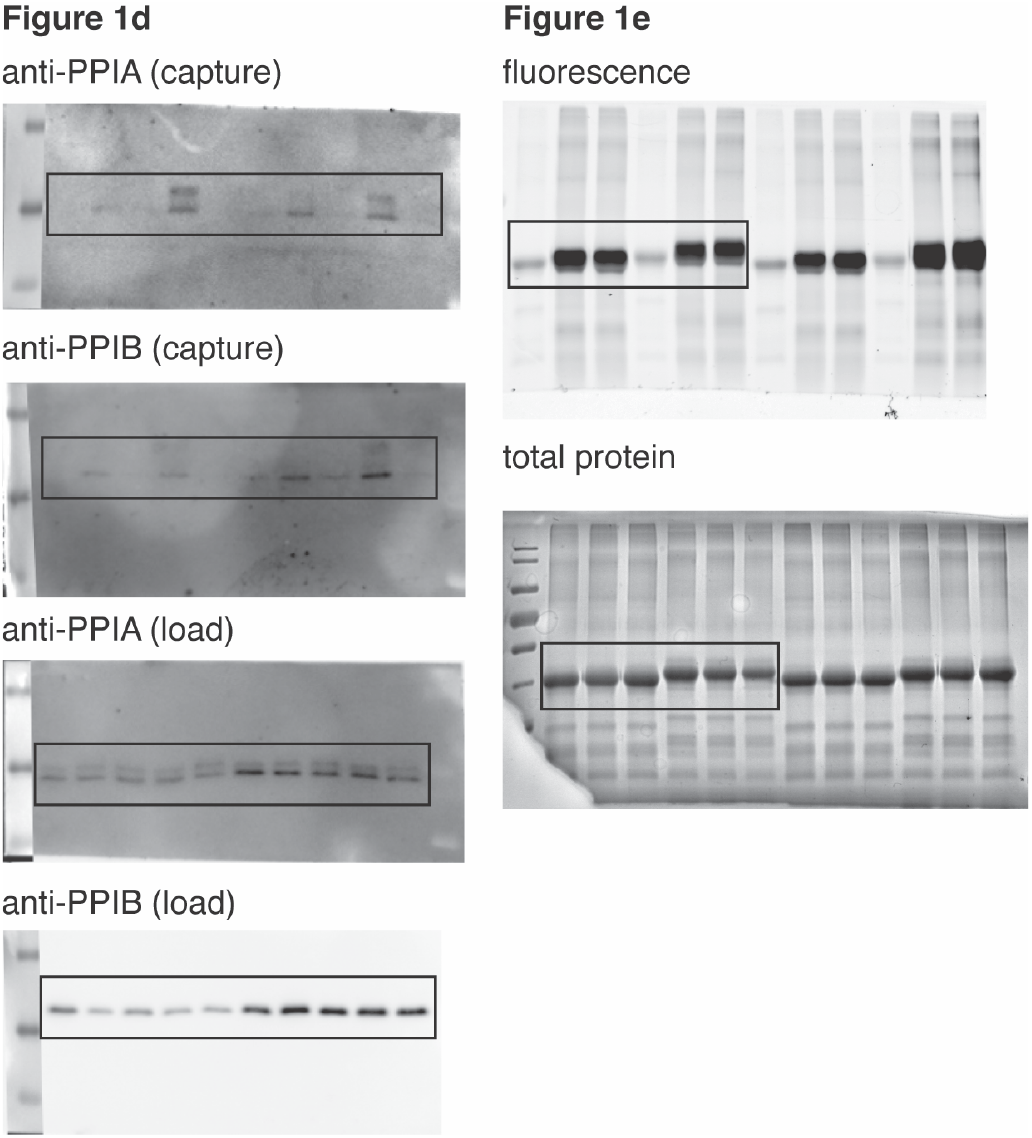
Uncropped Western Blots related to Figure 1.

**Supplementary Figure 2:**
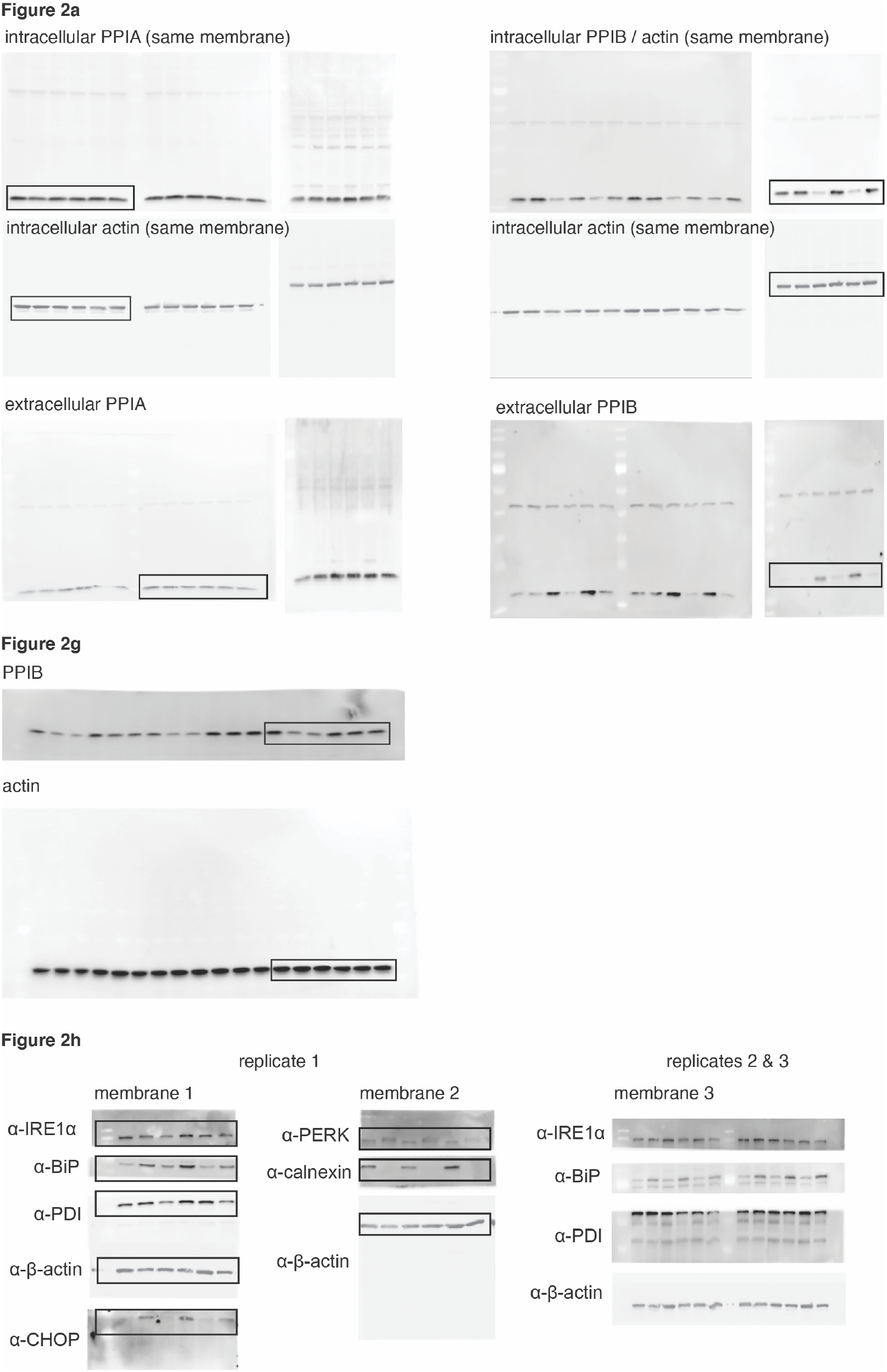
Uncropped Western Blots related to Figure 2.

**Supplementary Figure 3:**
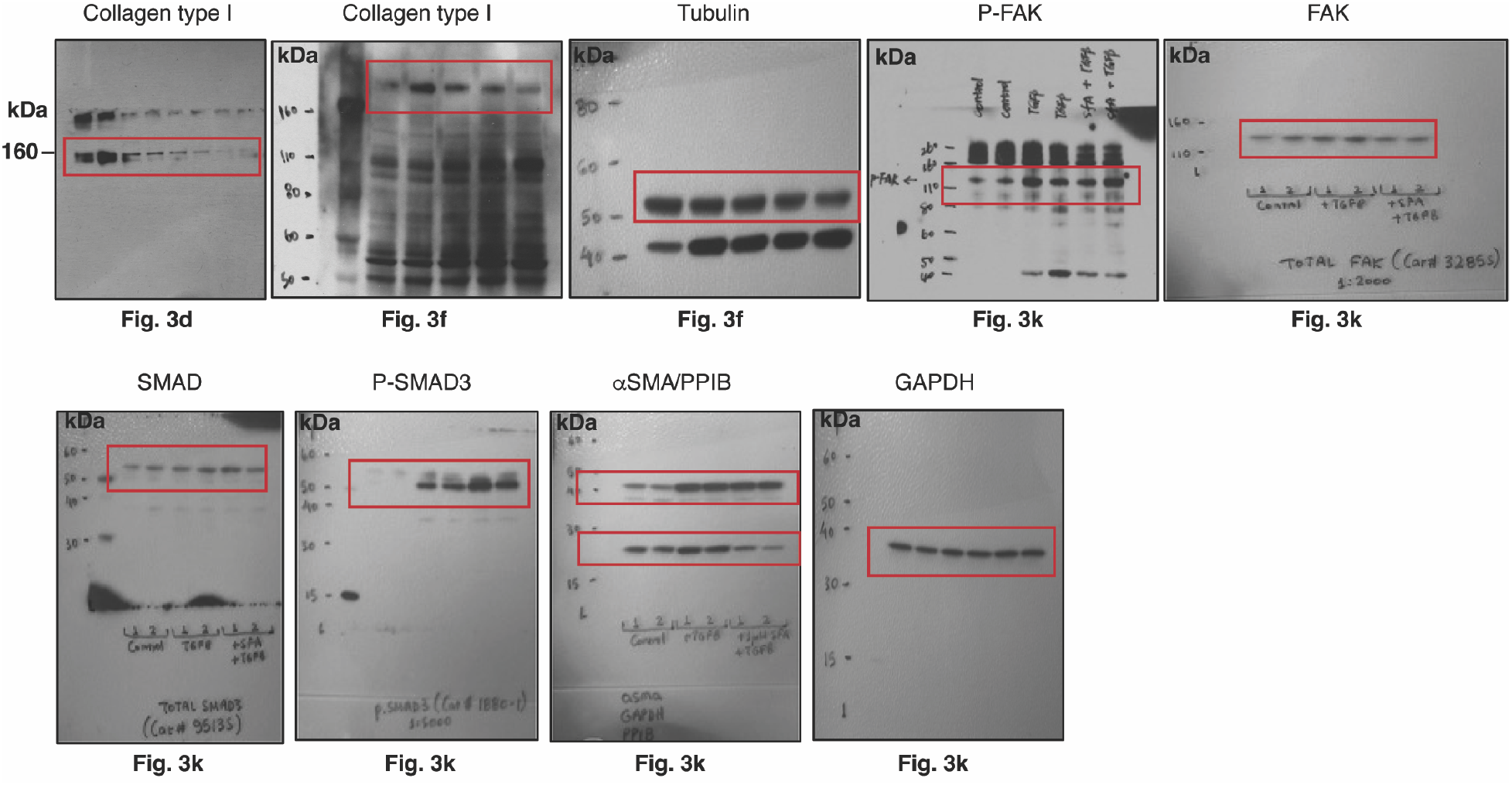
Uncropped Western Blots related to Figure 3.

**Supplementary Figure 4:**
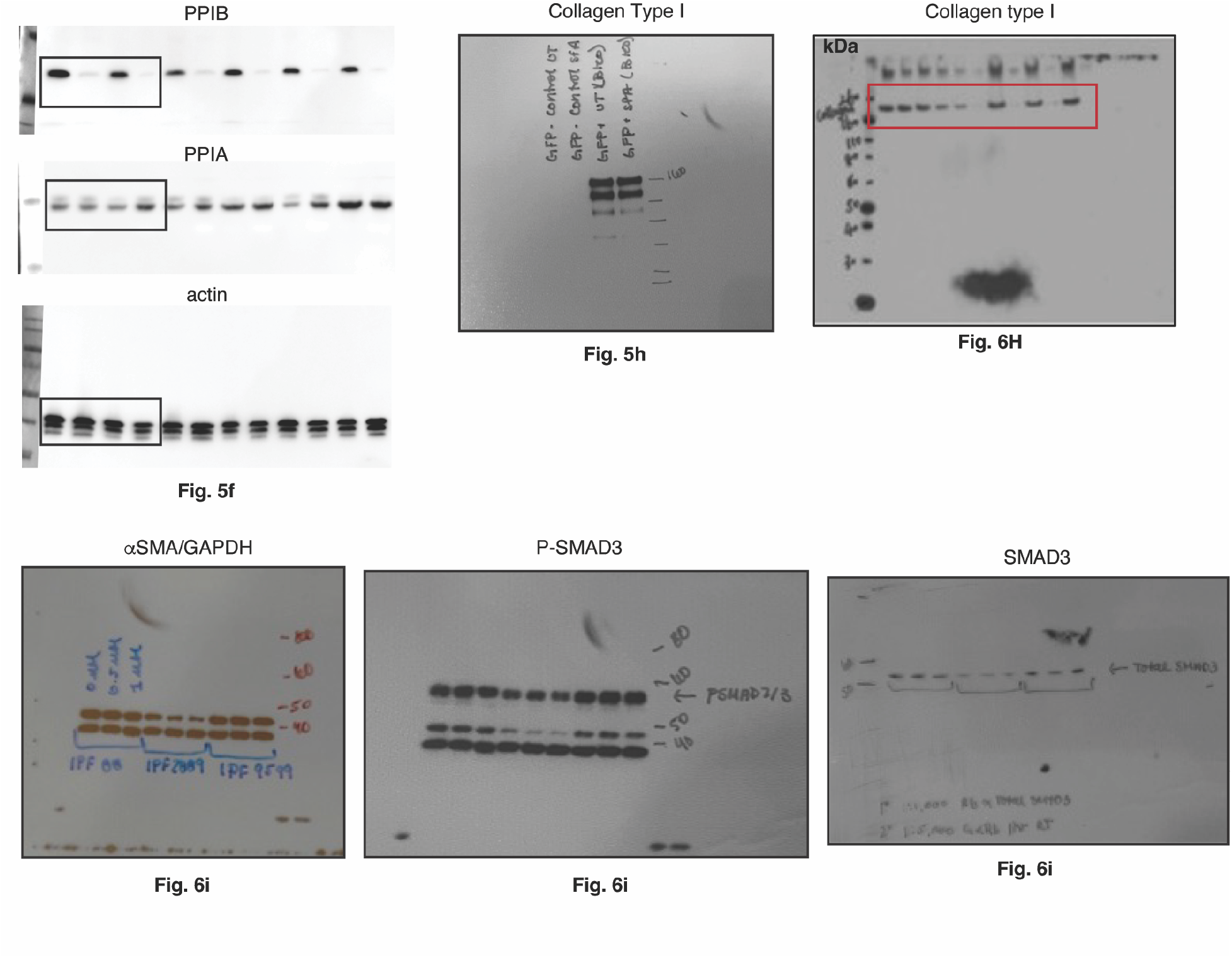
Uncropped Western Blots related to Figure 5, and Figure 6.

**Supplementary Figure 5:**
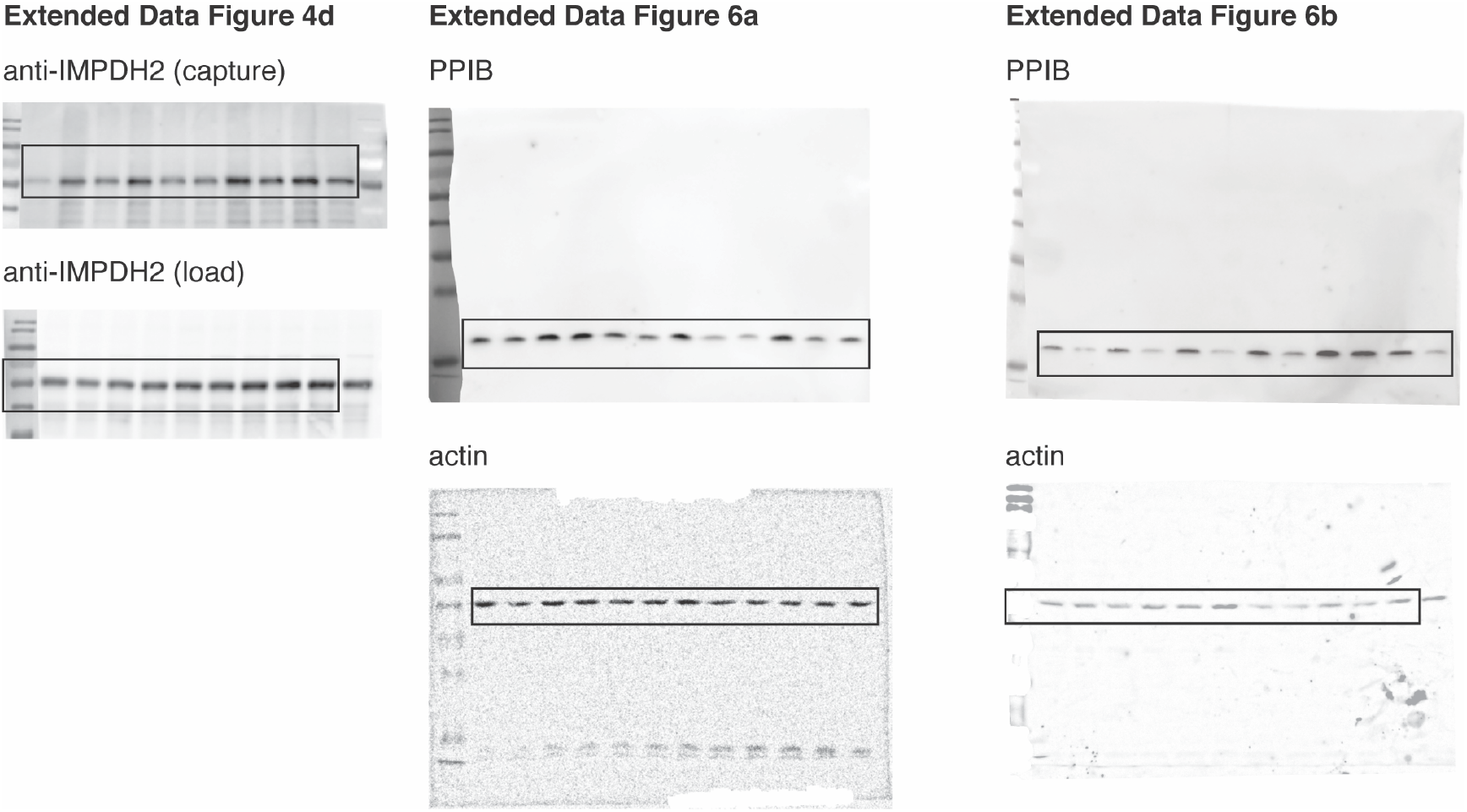
Uncropped Western Blots related to Extended Data Figure 4 and Extended Data Figure 6.

